# Nonsense-mediated decay controls a negative feedback loop in innate immune sensing

**DOI:** 10.1101/2025.05.09.652687

**Authors:** Simon Boudreault, Yahira Rivera-Lopez, Max B Ferretti, James Bonner, Bertram L Jacobs, Kristen W Lynch

## Abstract

Nonsense-mediated decay (NMD) is an mRNA decay pathway which degrades potential harmful transcripts that contain premature termination codons. However, NMD’s importance also extends to the control of isoform abundance under physiological conditions. During viral infection, NMD is inhibited through numerous mechanisms; however, NMD has been shown to have both antiviral as well as proviral activities, raising further questions into the role and control of NMD during viral infection. These observations have led us to investigate the potential involvement of NMD in dsRNA sensing as a mechanism that might explain these discrepancies. Using EIF4A2 exon 10B inclusion as an example of AS-NMD isoform accumulating during viral infection, we show that dsRNA sensing inhibits NMD. This effect is correlated with translational blockade and is driven primarily by RNaseL activation, and by PKR in the absence of RNaseL activation. Surprisingly, NMD inhibition limits the induction of IFN-β as well as interferon-stimulated genes, and this effect is upstream of IRF3 phosphorylation and translocation to the nucleus. NMD inhibition also decreases PKR and RNaseL activation as well as PIC-mediated cell death by decreasing the dsRNA content, suggesting NMD directly controls dsRNA sensing by controlling the dsRNA load. Therefore, inhibition of NMD upon dsRNA sensing provides a negative feedback loop that contributes to shaping the innate immune sensing pathways.

**SIGNIFICANCE:** Nonsense-mediated decay (NMD) is a translation-dependent mRNA decay pathway that plays an important role in shaping the transcriptome. In this manuscript, we show that dsRNA sensing, as is typical during viral infection, inhibits NMD mainly through the translational inhibition caused by RNaseL activation. This NMD inhibition forms a negative-feedback loop that limits dsRNA sensing, thus preventing overactivation of dsRNA-mediated pathways. These findings contribute to a better understanding of the molecular mechanisms that limit antiviral responses as well as inflammation and inform the critical role that mRNA processes plays in innate immunity.

## INTRODUCTION

Nonsense-mediated decay (NMD) is an eukaryotic translation-dependent mRNA decay pathway that detects and targets specific transcripts for degradation^1^. Activation of NMD requires the recruitment and phosphorylation of the UPF1 helicase by the SMG1 kinase, usually downstream of a translation termination codon, as the first step to commit the transcript to decay^2,3^. Decay can then occurs through two different pathways, either by endonucleolytic cleavage mediated by SMG6, or recruitment of the CCR4/NOT complex for deadenylation through SMG5 and SMG7^4^. A major feature that induces NMD is the presence of an exon-junction complex downstream of the termination codon, which typically indicates the presence of a premature stop codon. In addition, the length of the 3’-UTR, RNA structures, and binding of RNA-binding proteins all contribute to the sensitivity of a transcript to NMD. The complex interplay between all these factors ultimately dictates whether the transcript is targeted by NMD, and the extent to which decay occurs^5^.

It has been estimated that NMD controls the expression of roughly 10% of endogenous unmutated mammalian mRNA to fine-tune their levels to various physiological conditions^6^. One way in which unmutated transcripts are targeted by NMD is via alternative splicing in a process called AS-NMD, in which introduction or removal of coding regions by alternative splicing leads to a premature termination codon that is then sensed by the NMD machinery^7–9^. Thus, products of alternative splicing that do not encode a full-length protein are actively degraded to limit potential harmful effects. Although viruses have been shown to trigger widespread alternative splicing changes during infection, the extent to which these splicing events are targets of NMD has not been thoroughly explored^10–14^.

Despite a dearth in investigation of AS-NMD during viral infection, the general importance of NMD during viral infection has been clearly established in recent years. NMD usually exerts an antiviral effect on viruses, especially RNA viruses with a positive sense single-stranded genome (+ssRNA viruses), including flaviviruses and Semliki Forest virus^15–20^. Viral RNA from these viruses exhibits NMD-triggering features and are often subject to NMD degradation. Consequentially, numerous viruses have been shown to directly target the NMD machinery to inhibit this antiviral process. For example, the capsid protein of Semliki Forest virus interacts with NMD components and can inhibit NMD upon ectopic expression^21^; UPF1 is targeted by the nucleocapsid of SARS-CoV-2^22,23^; and the exon-junction complex protein PYM1 is targeted by flaviviruses capsid proteins to inhibit NMD^16^. However, PYM1 is also a proviral factor during another flavivirus infection, Hepatitis C virus (HCV), raising questions about the broader role of NMD during viral infection^24^. UPF1 is also proviral for human immunodeficiency virus (HIV)^25^, while negative-strand RNA viruses (−ssRNA viruses) such as Influenza A virus (IAV) and respiratory syncytial virus are not sensitive to the effects of NMD^26^. These conflicting results about the function of NMD during viral infections suggest a more intricate and nuanced role of NMD during virus-host interactions that requires further studies.

Virus-host interactions describe the complex relationship between the host cell and the invading pathogen, which ultimately dictates the outcome of infection. During their replication cycle, viruses produce a plethora of pathogen-associated molecular patterns (PAMPs), such as incompletely capped mRNA and dsRNA, that are sensed by various host receptors to mount an efficient antiviral response. Upon sensing of dsRNA, the response can be divide into three main arms: (1) activation of the interferon (IFN) response pathway through RIG-I and MDA-5, leading to the production and secretion of IFN-β that stimulates the induction of interferon-stimulated genes (ISG) that are mostly antiviral^27^; (2) activation of the OAS-RNaseL system, in which RNaseL cleaves mRNA, rRNA, as well as viral RNA to inhibits translation and impair viral replication^28^; and (3) activation of PKR, which phosphorylates EIF2α to impair translation^29^. The contribution of NMD to these responses to dsRNA has not been deciphered.

In the present study, we demonstrate a broad program of AS-NMD in response to both viral infection and dsRNA sensing. We then investigate the drivers and impact of NMD upon dsRNA sensing, using EIF4A2 exon 10B inclusion as an example of AS-NMD. We show that dsRNA sensing inhibits NMD, mainly through RNaseL activation, and NMD inhibition itself restricts dsRNA-mediated cellular processes, such as the IFN response and PKR activation. These data reveal a negative feedback loop involving NMD in dsRNA sensing that contributes to shaping the innate immune sensing pathways.

## RESULTS

### EIF4A2 exon 10B is representative of NMD-targeted isoforms accumulating during viral infections

A well-documented role of NMD in cells is to reduce the abundance of alternative spliced products that generate premature stop codons, in a process known as AS-NMD^7–9^. Given that viral infection has been reported to have a broad impact on alternative splicing^12,14^ and to inhibit NMD^17,18,20^, we first asked if some of the previously observed viral-induced alternative splicing changes might be attributable to accumulation of NMD-targeted isoforms. Indeed, we find that roughly 30% of alternative splicing changes triggered by Influenza A Virus (IAV)^13^, as well as triggered by Vesicular Stomatitis Virus (VSV), are also observed in response to NMD inhibition (**Figure 1A**). This significant proportion of alternative splicing changes being NMD-sensitive suggests that viral infection has a profound and broad effect on the NMD process, leading to an accumulation of spliced transcripts that would normally be removed by NMD.

**Figure 1.**
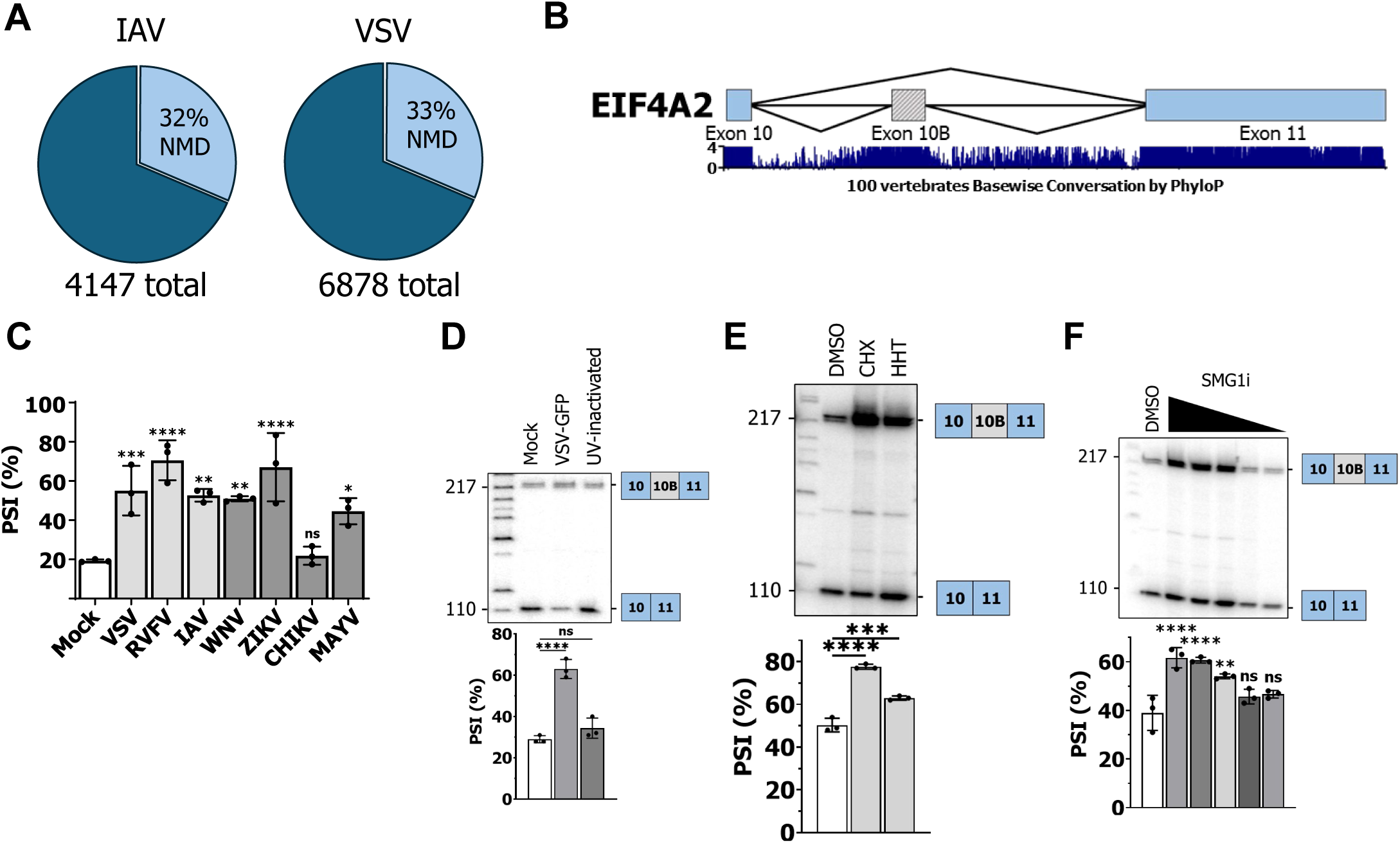
EIF4A2 exon 10B is representative of NMD-targeted isoforms accumulating during viral infections. (A) Widespread accumulation of NMD isoforms upon infection with Influenza A virus (IAV) and Vesicular Stomatitis Virus (VSV). A549 cells were infected for 12 h (IAV) or 24 h (VSV), RNA sequenced, splicing analyzed using MAJIQ and NMD isoforms determined from NMD datasets (see material and methods). (B) Schematic of the exon 10B splicing in EIF4A2. Conservation values across 100 vertebrates is shown underneath. (C) Impact of viral infection with prototypical -ssRNA viruses (Vesicular Stomatitis Virus (VSV), Rift Valley Fever Virus (RVFV), and Influenza A Virus (IAV)) and +ssRNA viruses (West Nile Virus (WNV), Zika virus (ZIKV), Chikungunya virus (CHIKV), and Mayaro virus (MAYV) on EIF4A2 exon 10B inclusion. A549 cells were infected at a MOI of 5, RNA harvested at 24 h PI and subjected to radiolabelled RT-PCR for EIF4A2 exon 10B inclusion. (D) Impact of VSV and UV-inactivated VSV on EIF4A2 exon 10B splicing. (E) EIF4A2 exon 10B-containing transcripts accumulate following translation inhibition. A549 cells were treated with DMSO, cycloheximide (CHX), or homoharringtonine (HHT) for 5h. RNA was harvested and subjected to radiolabelled RT-PCR for EIF4A2 exon 10B inclusion. (F) EIF4A2 exon 10B-containing transcripts accumulate following NMD inhibition. A549 cells were treated with DMSO or SMG1i, a SMG1i inhibitor, at 3 μM, 1 μM, 300 nM, 100 nM, or 30 nM, for 7h. RNA was harvested and subjected to radiolabelled RT-PCR for EIF4A2 exon 10B inclusion.

To further investigate the phenomena of AS-NMD in viral infection, we selected two previously characterized examples of AS-NMD, inclusion of alternative exon 10B in EIF4A2, and a differential 5’-SS usage in CDKN2AIP (**Figure 1B and Supplementary Figure S1A**). Both of these AS-NMD isoforms have previously been described to be induced upon infection with reovirus and IAV^10,11,13^ and EIF4A2 exon 10B accumulation was also recently observed during coronavirus infection^30^. To expand these previous results, we quantified splicing in A549 cells infected with a broad range of RNA viruses encompassing prototypical negative-strand RNA viruses (−ssRNA viruses; Vesicular Stomatitis Virus; VSV, a vesiculovirus, Rift Valley Fever Virus; RVFV, a phlebovirus; and Influenza A Virus, IAV, an orthomyxovirus), and positive-strand RNA viruses (+ssRNA viruses; West Nile Virus; WNV and Zika virus; ZIKV, two flaviviruses, and Chikungunya virus; CHIKV and Mayaro virus; MAYV, two togaviruses). RNA was harvested 24 h post-infection and subjected to RT-PCR to monitor the inclusion of EIF4A2 exon 10B and the alternate 5’-SS use in CDKN2AIP. All of the viruses tested, with the partial exception of CHIKV, showed a significant increase in exon 10B inclusion following infection and differential 5’-SS usage in CDKN2AIP (**Figure 1, Supplementary Figure S1B**). The percent spliced-in (PSI) metric was used to quantitate the level of inclusion in these alternative splicing events; it represents the percentage of the long form (10B included) over total abundance (both 10B included and 10B excluded). Viral replication was required to induce these AS-NMD isoforms, as UV-inactivated viruses were unable to trigger changes in isoform expression of EIF4A2 or CDKN2AIP (**Figure 1D, Supplementary Figure S1B**). Increased 10B inclusion could also be observed in VSV-infected HEK 293T or murine L929 fibroblasts, arguing against a cell-type specific effect observed only in A549 (**Supplementary Figure S2**).

To additionally confirm that 10B inclusion is normally repressed by NMD in our system, we first monitored the responsiveness of 10B to translation inhibition, since active translation is required for NMD^1^. Consistently, treatment of A549 cells with the translation inhibitors cycloheximide or homoharringtonine triggered 10B accumulation (**Figure 1E**). As further demonstration that inhibition of NMD leads to increased abundance of EIF4A2 exon 10B inclusion, we treated A549 cells with an inhibitor of SMG1 (SMG1i), the kinase responsible for phosphorylation of UPF1 as the first step for NMD^31^. Treatment with SMG1i was confirmed to decrease the phosphorylation of UPF1, a widely used readout for the level of steady-state decay by NMD^2^ (**Supplementary Figure S1C**). Upon treatment with increasing SMG1i concentrations, we observe a dose-dependent increase in the accumulation of 10B-containing isoforms, confirming that 10B-containing transcripts are normally reduced by NMD in A549 cells (**Figure 1F**). Similar results were also obtained with the 5’-SS in CDKN2AIP (**Supplementary Figure S1D**). In conclusion, inclusion of EIF4A2 exon 10B or altered use of the regulated 5’-SS in CDKN2AIP direct the resulting transcripts for NMD degradation. As such, both of these splicing events are representative NMD isoforms increasing following infection with numerous RNA viruses. As inclusion of EIF4A2 exon 10B is the most straightforward to quantify by RT-PCR, we use this isoform expression as a marker for NMD in our subsequent experiments.

### DsRNA sensing increases EIF4A2 exon 10B inclusion by inhibiting NMD

The observation that changes in splicing associated with AS-NMD are triggered during infection by numerous different viruses, and that this phenomenon is dependent on viral replication, suggests that all these viruses produce something that blocks NMD. Numerous viruses produce pathogen-associated molecular patterns (PAMP) such as dsRNA and 5’ triphosphate RNA during replication that are sensed by the host cell^32^. To test if a PAMP might be at least one mechanism responsible for the inhibition of NMD, we transfected seven different commercially available PAMPs in A549 and monitored their effect on NMD by assaying EIF4A2 exon 10B inclusion. Of all the PAMPs tested, only poly(I:C) (PIC), a mimic of dsRNA, could robustly trigger increased 10B inclusion in A549, as well as in HEK 293T or L929 **(Figure 2A, Supplementary Figure S3**). A block in NMD by PIC treatment is consistent with reports that dsRNA sensing inhibits translation^29^, which we confirmed in our system by puromycin-incorporation (**Figure 2B**). To further confirm that the effect of PIC on EIF4A2 exon 10B inclusion is attributable to inhibition of NMD, we monitored the level of phosphorylated UPF1 (pUPF1). As the first step of NMD is the phosphorylation of UPF1 by the SMG1 kinase, pUPF1 is a widely used readout for the level of steady-state activity of NMD^2^. Upon PIC transfection, we observed a 50% decrease in pUFP1, most readily observed upon accumulation of pUFP1 using the phosphatase inhibitors okadaic acid and calyculin^33^ (**Figure 2C**). Finally, we analyzed publicly available RNA-Seq data from PIC-treated A549 cells and again find a high level of AS-NMD isoforms induced in response to PIC (**Figure 2D**). Together, these results show that upon sensing of dsRNA, decreased efficiency of translation and NMD correlates with an increase in abundance of NMD-targeted isoforms.

**Figure 2.**
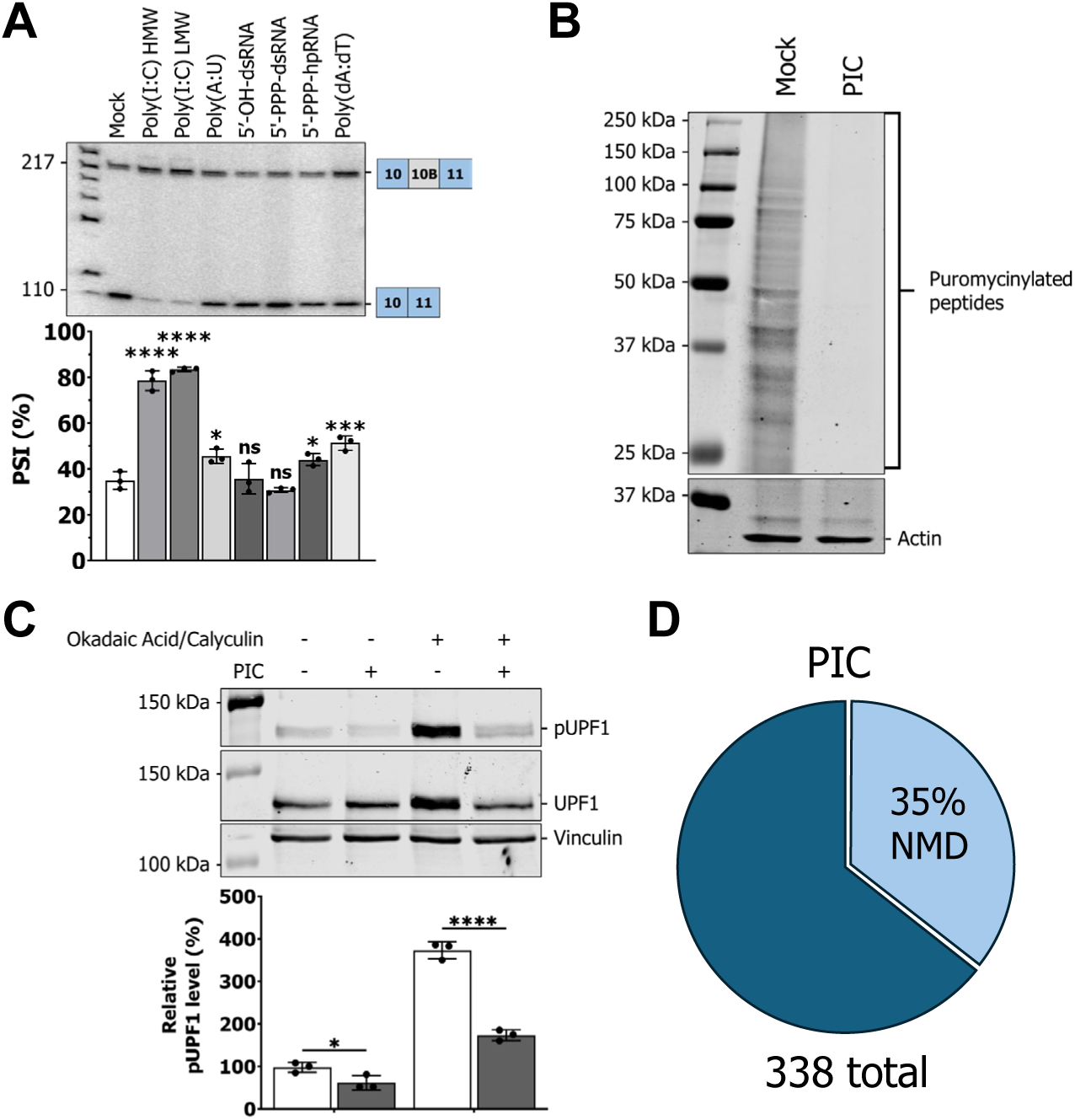
DsRNA sensing increases EIF4A2 exon 10B inclusion by inhibiting NMD. (A) Monitoring of EIF4A2 exon 10B inclusion following transfection of seven different PAMP in A549 cells for 6h. (B) Puromycin-incorporation assay of mock-transfected or PIC-transfected A549 cells. Cells were treated with 10 µg/mL of puromycin for 10 min, prior to western-blotting for puromycinylated peptides or actin loading control. (C) Western blot of pUPF1, UPF1, and vinculin (loading control) in A549 cells following PIC transfection. 50 nM calyculin and 500 nM okadaic acid were added 5 minutes prior to harvesting to prevent dephospholyration of UPF1 and increase the signal, where indicated. (D) Widespread accumulation of NMD isoforms upon PIC transfection. Publicly available datasets of PIC-transfected cells were analyzed using MAJIQ for alternative splicing and NMD isoforms determined from NMD datasets (see material and methods).

### NMD inhibition during dsRNA sensing is mediated primarily by RNaseL, and, to a lesser extent, PKR

Having established in our system that dsRNA sensing leads to a blockade in NMD, we next sought to decipher the precise sensors responsible for this NMD inhibition. Since we hypothesize that this block in NMD is downstream of translational inhibition, we turned our attention to two of the main factors controlling translation during dsRNA sensing, i.e. RNaseL and PKR. Upon binding to dsRNA, 2’-5’-oligoadenylate synthetases (OAS1, OAS2, and OAS3) catalyze the production of 2’-5’-oligoadenylates that trigger the dimerization of RNaseL, a nonspecific RNase that cleaves rRNA, mRNA, and viral RNA. This widespread RNA cleavage by RNaseL hampers translation^28^. By contrast, PKR is directly activated by dsRNA and phosphorylates the translation initiation factor eIF2α, which potently inhibits translation^29^.

We tested OAS1, OAS2, OAS3, RNaseL, and PKR A549 knockout (KO) cells for their capacity to increase EIF4A2 exon 10B inclusion upon PIC transfection (**Figure 3A**). In A549 cells, RNaseL activation has been demonstrated to be solely dependent on OAS3^34^, which we also confirmed (**Supplementary Figure S4**). Accordingly, genetic ablation of RNaseL and OAS3 both lead to a roughly 50% reduction in the increase in exon 10B inclusion, suggesting that RNaseL activation at least partially mediates NMD inhibition, while knock out of OAS1 and OAS2 had no effect. Notably, PKR knockout also did not reduce the ability of PIC to trigger increased exon 10B inclusion. However, given the fact that RNaseL depletion only has a partial impact on exon 10B inclusion, we wondered if PKR could be compensating in the absence of RNaseL. Indeed, depletion of PKR using siRNA in OAS3 KO (**Figure 3B**, left) or RNaseL KO (**Figure 3B**, right) cells completely blocks any PIC-induced inclusion of exon 10B; PKR depletion using siRNA was confirmed by WB (**Supplementary Figure S5A).** We also assessed the impact of the depletion using siRNA of RNaseL in the A549 PKR KO cells; although the siRNA-mediated depletion of RNaseL does not completely abrogate RNaseL activity, in the absence of PKR a significant reduction of the impact on 10B could be observed (**Supplementary Figure S5A, B**). Together, these data suggest that RNaseL is the dominant driver of PIC-induced inhibition of AS-NMD, with PKR activity compensating for this effect in the absence of RNaseL activation.

**Figure 3.**
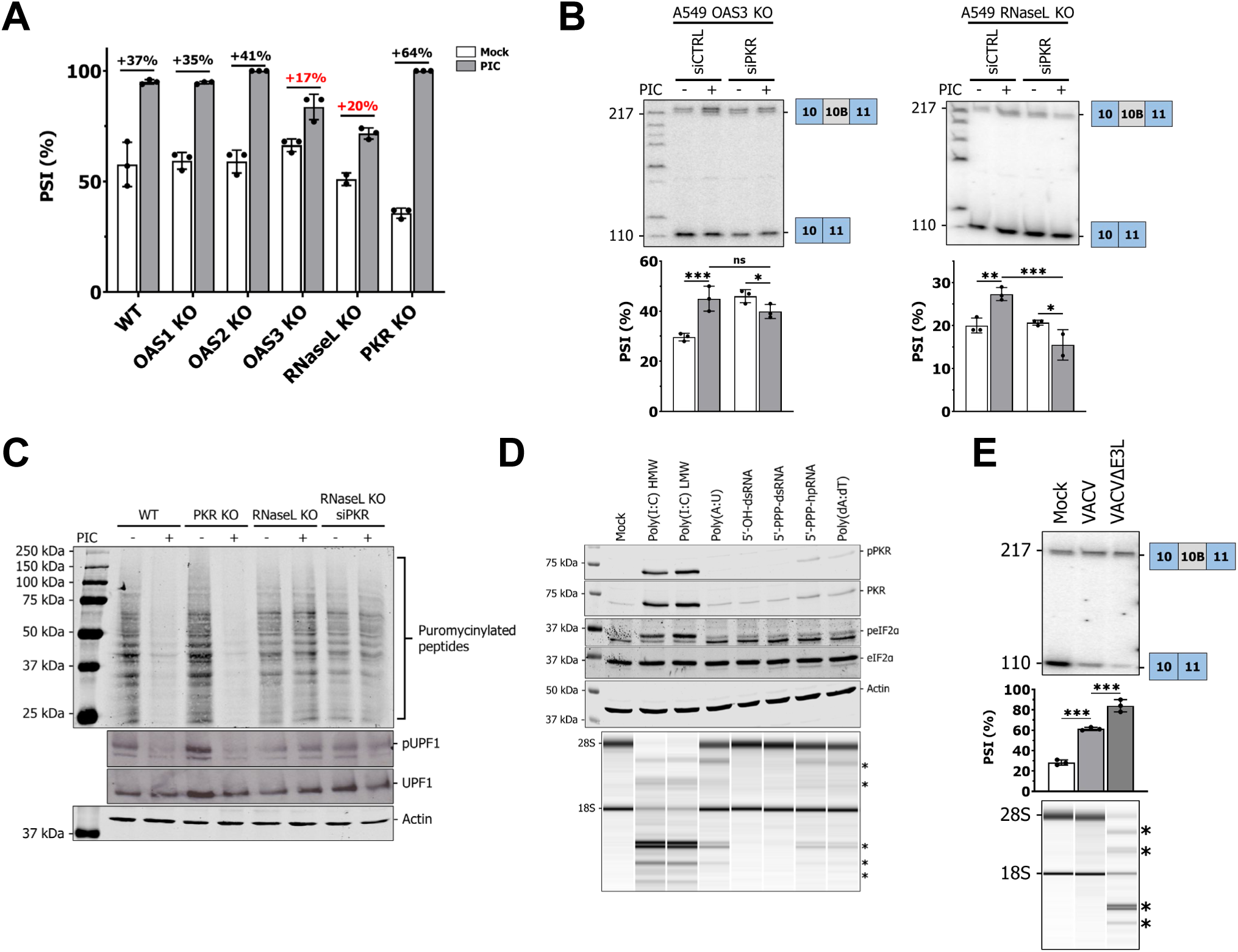
NMD inhibition during dsRNA sensing is mediated primarily by RNaseL, and, to a lesser extent, PKR. (A) Impact of PIC transfection on 10B inclusion in WT, OAS1, OAS2, OAS3, RNaseL, and PKR KO A549 cells. (B) siRNA-mediated depletion of PKR in OAS3 or RNaseL KO A549 cells. Cells were transfected with siRNA for 72h prior to be transfected with PIC for 6h and RNA harvested for EIF4A2 exon 10B radioactive RT-PCR. Two-way ANOVA with uncorrected Fisher’s LSD multiple comparisons test. (C) Translational output and NMD status in WT, PKR KO, RNaseL KO, and RNaseL KO + siPKR upon stimulation with PIC. Top, puromycin incorporation assay to measure translation; bottom, WB against pUPF1; cell were treated with 50 nM calyculin and 500 nM okadaic acid 5 minutes prior to harvesting to increase pUPF1 signal. (D) Monitoring PKR and RNaseL activation following PAMP transfection in A549. Top, western-blot for pPKR, total PKR, peIF2α, total eIF2α as well as actin (loading control). Bottom, total RNA was run on a Bioanalyzer to monitor RNaseL activation by monitoring rRNA cleavage. Asterisks denote the cleavage products of ribosomal RNA typical of RNaseL activation. (E) Monitoring WT vaccinia virus (VACV) or ΔE3L mutant (VACVΔE3L) impact on 10B inclusion. Top, radioactive RT-PCR for EIF4A2 exon 10B; bottom, total RNA on a Bioanalyzer to monitor RNaseL activation. Asterisks denote the cleavage products of ribosomal RNA typical of RNaseL activation. Ordinary one-way ANOVA using Tukey’s multiple comparisons test.

Although EIF4A2 exon 10B inclusion is a reliable marker for NMD, we further confirmed that RNaseL and PKR block translation, which is required for NMD. Indeed, RNaseL is responsible for the bulk of the translational blockade upon PIC transfection, mirroring its effect on EIF4A2 exon 10B inclusion (**Figure 3C**). Moreover, RNaseL is also the main driver for reducing pUPF1 during dsRNA sensing, further confirming its primary role in controlling the NMD status upon recognition of dsRNA (**Figure 3C**). Additionally, we find that activation of RNaseL, as well as PKR and eIF2α phosphorylation correlates with the degree of impact on exon 10B inclusion in response to PAMP (**Figure 3D**). Specifically, we find that only PIC triggers strong RNaseL/PKR activation, with modest impact on RNaseL by Poly(A:U), 5’-PPP-hpRNA, as well as Poly(dA:dT) (**Figure 3D**).

Finally, to directly determine if dsRNA sensing during infection mediates the effect on 10B inclusion, we turned to vaccinia virus, a dsRNA virus that replicates solely in the cytoplasm. Removal of E3L, a viral dsRNA/dsDNA binding protein, leads to enhanced dsRNA sensing during infection as well as increased RNaseL activation^34^. We monitored exon 10B inclusion upon infection with WT as well as ΔE3L vaccinia virus and shown that 10B inclusion is significantly increased in the ΔE3L mutant together with strong RNaseL activation (**Figure 3E**). Altogether, these results support a model in which translational inhibition, driven mainly by RNaseL, and to some extend by PKR in the absence of RNaseL activation, are responsible for the accumulation of NMD isoforms upon dsRNA sensing.

### NMD controls the IFN response

The observation that NMD is inhibited during dsRNA sensing led us to question if this inhibition could be playing a functional role in the events that are triggered following dsRNA recognition and signalling. We first tested the impact of NMD inhibition on the early events leading to the production of type I interferons by stimulating cells with PIC in the presence of SMG1i and observed no impact of SMG1i on the PIC-induced expression of IFN-β or the downstream interferon-stimulated genes (ISG) IFIT1/2 (**Supplemental Figure S6A**).

One possible cause of the lack of impact of SMG1i on the interferon pathway is our observation that NMD is strongly inhibited following PIC transfection (**Figure 2A-C**); we thus wondered if the NMD inhibition already induced during PIC infection could mask the effect of SMG1i. To test this hypothesis, we leveraged our finding that PIC-induced inhibition of NMD is driven mainly by RNaseL, and to a lesser extent, by PKR (**Figure 3A, B**). We thus transfected WT, PKR KO, RNaseL KO, or RNaseL/PKR double-depleted cells with PIC, and assessed induction of IFN-β, IFIT1, and IFIT2 by qPCR. Notably, compared to wild type cells, we observed a significant increase in the IFN-β and ISG induction in RNaseL KO, and RNaseL/PKR double-depleted cells, but not in PKR KO cells (**Figure 4A, white bars**). Moreover, treatment of the RNaseL KO or RNaseL/PKR double-depleted cells with SMG1i significantly reduced IFN-β and ISG induction to a degree correlated with the amount of NMD (**Figure 4A, grey bars**). These results support a model in which NMD inhibition during dsRNA sensing decreases the IFN response, and abrogation of the NMD inhibition by removal of RNaseL and PKR relives this restriction. The conclusion that NMD inhibition limits the interferon response is further supported by the fact that ISG expression as well as IFN-β induction can be reduced by SMG1i in cellular context where NMD inhibition is relieved upon PIC transfection (RNaseL KO or RNaseL/PKR double-depleted cells). In addition, we validated that NMD inhibition has only a marginal effect on direct ISGs induction in the absence of dsRNA sensing, by testing the impact of SMG1i on ISGs expression in cells treated with different IFN-β concentrations (**Supplemental Figure S6B**). Therefore, in all of these experiments, the degree of NMD inhibition is strongly correlated with the extent of the interferon response, and NMD activity is required for optimal IFN response induction at the IFN-β induction step.

**Figure 4.**
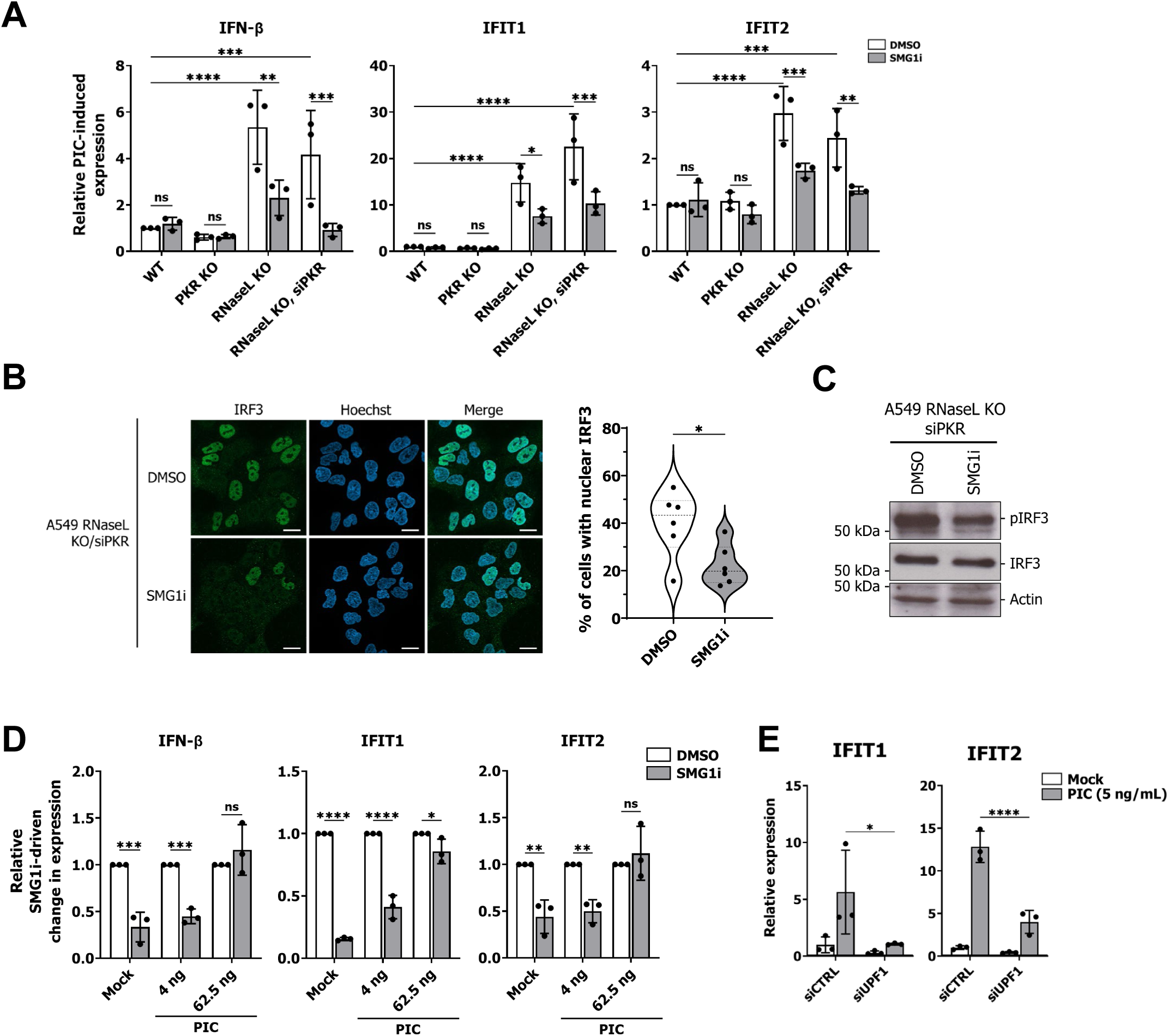
NMD controls the IFN response. (A) A549 WT, PKR KO, RNaseL KO as well as RNaseL KO transfected with siRNA against PKR were pre-treated with DMSO or SMG1i (3 μM) for 1h and transfected with 1 μg/mL of PIC and RNA harvested at 6h. Two-way ANOVA with uncorrected Fisher’s LSD multiple comparisons test. (B) Left, immunofluorescence for IRF3 of DMSO or SMG1i-treated cells transfected with PIC. A549 RNaseL KO were transfected with siPKR for 72h, pre-treated with DMSO or SMG1i (3 μM) for 1h and transfected with 1 μg of PIC prior to IF. Scale bar, 20 μm. Right, quantification of the number of cells with IRF3 nuclear localization. 6 fields were counted for each condition each containing between 14 and 36 cells. Unpaired two-tailed Student’s t-test. (C) WB against pIRF3 in RNaseL KO siPKR cells. Cells were pre-treated with DMSO or SMG1i (3 μM) for 1h and transfected with 1 μg/mL of PIC and proteins harvested after 6h. (D) A549 cells were pre-treated with SMG1i (3 μM) or DMSO for 1h, transfected with 0, 4, or 62.5 ng/mL of PIC for 6h prior to RNA harvesting and qPCR for IFN-β and IFIT1/2. DMSO control for each biological replicate was normalized to 1 to isolate the effect of SMG1i. Two-way ANOVA with Šídák’s multiple comparisons test. (E) A549 were transfected with control siRNA or siRNA against UPF1 for 72h prior to transfection with no PIC (mock) or 5 ng/mL of PIC, RNA harvested at 6h, and submitted to qPCR for IFIT1 and IFIT2. Two-way ANOVA with Šídák’s multiple comparisons test.

We next sought to understand how NMD is exerting control on the IFN response. Since we observed concurrent decreases in IFN-β induction with ISG induction, we hypothesize that the effect of NMD is upstream of IFN-β induction. We thus monitored the impact of NMD on translocation of IRF3 in the nucleus, which is a driving step in IFN-β mRNA production. To test this, we assessed the impact of SMG1i on IRF3 translocation in WT and RNaseL/PKR double-depleted A549 cells stimulated with PIC (**Figure 4B, Supplemental Figure S7**). Upon treatment with SMG1i, we observed a significant reduction in the percentage of RNaseL/PKR double-depleted A549 cells with nuclear IRF3, suggesting NMD inhibition acts upstream of the translocation of IRF3 in the nucleus **(Figure 4B)**. Concurrent with the absence of effect on the IFN-β mRNA induction of SMG1i in WT cells (**Figure 4A**, **Supplemental Figure S6A**), we didn’t observe a statistically significant decrease in nuclear IRF3 in WT cells upon SMG1i treatment (**Supplemental Figure S7**). We also confirmed that the phosphorylation of IRF3 is reduced in RNaseL/PKR double-depleted cells upon NMD inhibition (**Figure 4C**).

We next wondered if we could establish this NMD-mediated negative regulation of the IFN response in WT cells. We reasoned that usage of less PIC could probably alleviate some of the effect on NMD without completely abrogating the IFN response and would provide a window in which to test the impact of NMD in a WT context using SMG1i. We first assessed NMD using EIF4A2 exon 10B inclusion across a broad range of PIC concentrations and indeed found that PIC concentrations of less than 250 ng/mL exhibit reduced EIF4A2 exon 10B inclusion (**Supplemental Figure S8**). Once again, the impact of PIC on exon 10B inclusion was correlated with the level of RNaseL activation (**Supplemental Figure S8**). We then stimulated wild type A549 with lower PIC concentrations (4 and 62.5 ng/mL), monitored the IFN response, and observed a reduction of IFN-β and ISG induction in the presence of SMG1i at 4 ng/mL that is abrogated at 62.5 ng/mL (**Figure 4D**). Importantly, these results, as well as the one from **Figure 4A**, could be replicated by using an siRNA against UPF1 instead of SMG1i, confirming that this effect is dependent on NMD inhibition, and not an off-target effect of SMG1i (**Figure 4E**, **Supplemental Figure S9**). Altogether, these results demonstrate that during dsRNA sensing, the inhibition of NMD decreases IRF3 phosphorylation and translocation to the nucleus, contributing to limiting IFN-β induction, and this effect is upstream of IRF3 phosphorylation/translocation to the nucleus.

### NMD controls dsRNA sensing

Since we observed NMD to be acting upstream of the phosphorylation and translocation of IRF3, we wondered if NMD could also be contributing more broadly to dsRNA sensing. In addition to the IFN response, dsRNA also activates PKR and RNaseL as shown in **Figure 3D**. We first monitored PKR activation following transfection with different PIC concentrations (**Figure 5A**). Although the lowest PIC concentration failed to activate PKR, at both 62.5 ng/mL and 1 μg/mL of PIC, PKR activation is reduced by SMG1i treatment, suggesting NMD also controls PKR activation in addition to the IFN response. We then monitored RNaseL activation following PIC transfection across a broad range of concentrations, in the presence or absence of the SMG1i. Importantly, at PIC concentrations below 250 ng/mL that don’t fully inhibit NMD, we do observe that further NMD inhibition with SMG1i reduces RNaseL activation by 30-40% (**Supplemental Figure S10**). Lastly, we wondered if NMD inhibition could provide protection against PIC-mediated cell death. WT or RNaseL/PKR double-depleted cells were transfected with PIC and the number of Sytox-orange positive cells (dead cells) were monitored using live-cell imaging (**Figure 5B**). In both cellular contexts, the cell death kinetics and the number of dead cells are drastically reduced by SMG1i, suggesting it also protects the cells from PIC-mediated cell death. Similar results could be observed using UPF1 depletion instead of SMG1i in WT A549 (**Supplemental Figure S10**).

**Figure 5.**
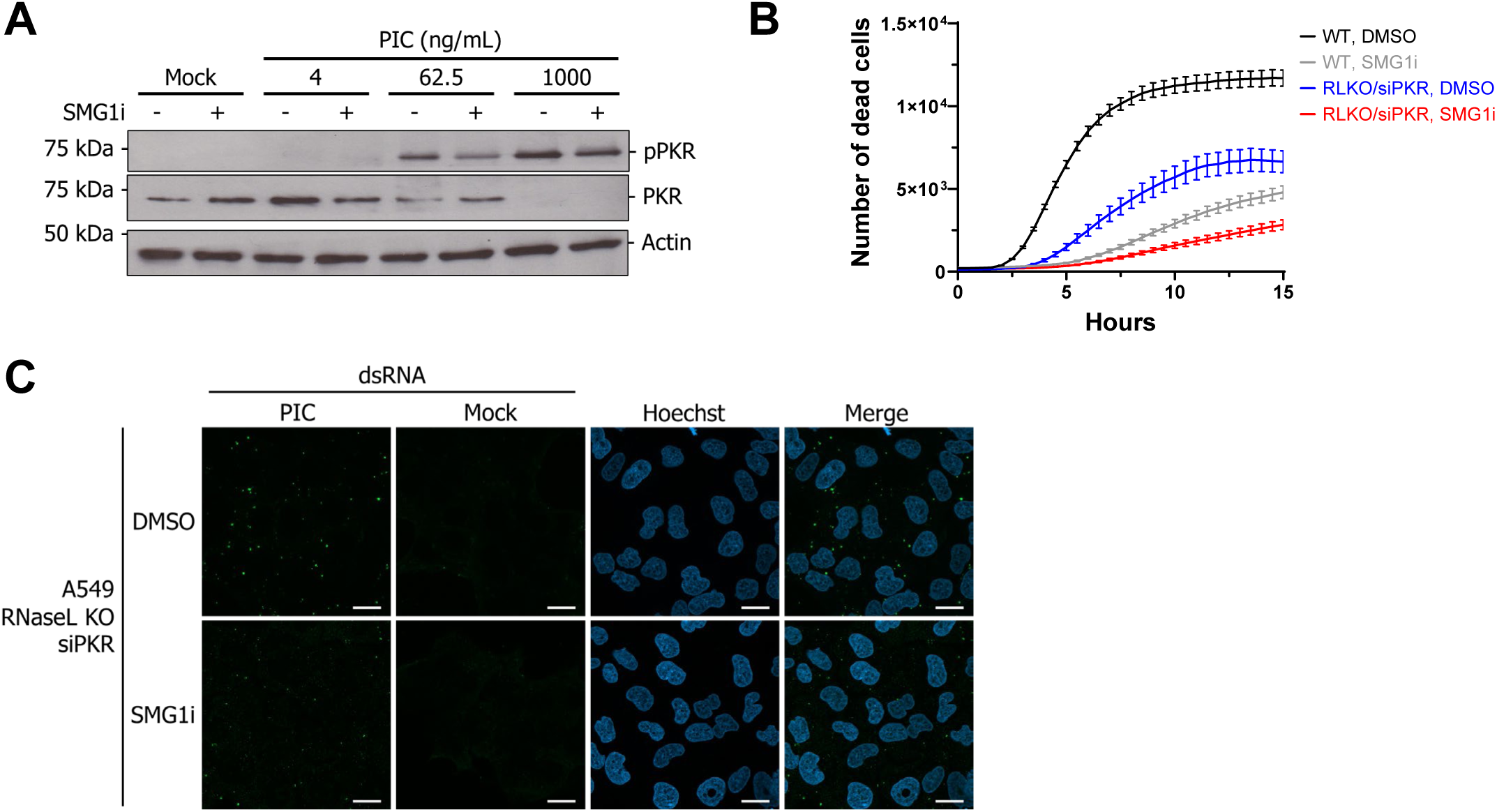
NMD controls dsRNA sensing. (A) WB to monitor the impact of SMG1i on PKR activation upon stimulation with 4, 62.5, and 1000 ng/mL of PIC. Cells were pre-treated with DMSO or SMG1i (3 μM) for 1h before PIC transfection with the indiquated concentraions of PIC; proteins were harvested 6h after stimulation. (B) Live-cell imaging of WT or RNaseL KO/siPKR A549 cells upon transfection of 1 μg/mL of PIC in the presence or absence of 3 μM of SMG1i. The number of dead cells correspond to Sytox-Orange positive cells. (C) Immunofluorescence for dsRNA using 9D5 of DMSO or SMG1i-treated cells transfected with PIC. A549 RNaseL KO were transfected with siPKR for 72h, pre-treated with DMSO or SMG1i (3 μM) for 1h and transfected with 1 μg/mL of PIC prior to IF. Scale bar, 20 μm.

Altogether, these results suggest that numerous dsRNA-mediated mechanisms – the IFN response, PKR activation, RNaseL activation, and PIC-mediated cell death – are decreased upon NMD inhibition and point to the effect of NMD lying at the dsRNA sensing step directly, either by masking dsRNA, or actively decreasing the dsRNA load to limit the activation of downstream pathways. To validate this hypothesis, we monitored the dsRNA content of PIC-transfected RNaseL/PKR double-depleted cells in the presence or absence of SMG1i (**Figure 5C**). Using the 9D5 antibody against dsRNA, we observe large, bright foci of dsRNA in DMSO-treated cells; upon SMG1i treatment, these foci are smaller, and the dsRNA signal is more diffuse. By contrast, in WT cells no obvious difference could be observed upon SMG1i treatment (**Supplemental Figure S11**). To further confirm the immunostimulatory potential of PIC in DMSO/SMG1i-treated A549, RNA harvested from these experiments was transfected again in A549 and the induction of IFIT1 measured (**Supplemental Figure S11**). We found that upon SMG1i treatment in RNaseL/PKR double-depleted cells, the RNA content containing the initial PIC transfected was less immunostimulatory, further supporting that the dsRNA content is reduced by NMD inhibition. These results underscore that in addition to controlling the IFN response, NMD also controls the other dsRNA-mediated processes such as PKR activation, RNaseL activation, and PIC-mediated cell death, and establish that the effect of NMD on dsRNA-mediated processes lies at the sensing step by reducing dsRNA load and sensing. Investigations into helicases that might be predicted to mediate the reduction of dsRNA sensing upon NMD inhibition, such as SKIV2L, DHX9, and EIF4A1/EIF4A2, have shown that none of these helicases are required for the impact of NMD inhibtion (**Supplemental Figure S10**). Therefore, future studies will be required to determine the mechanism(s) by which NMD, and its inhibition, impact dsRNA sensing in cells.

## DISCUSSION

The conflicting roles of NMD during viral infection, presenting both proviral and antiviral effects^17–20^, suggest that NMD has a deeper impact beyond our current understanding. Here we demonstrate, using an AS-NMD isoform in EIF4A2 as a readout for NMD, that dsRNA sensing inhibits NMD primarily through RNaseL activation, and to a lesser extent through redundant activity of PKR. Conversely, we also show that NMD itself is important for dsRNA-mediated processes, as its inhibition decreases dsRNA sensing and subsequent triggering of the IFN response, as well as activation of PKR, RNaseL, and PIC-mediated cell death.

Inhibition of NMD has been widely reported to be induced upon viral infection^17–20^. The production of dsRNA by viruses^35,36^, as well as translation inhibition triggered during their replication cycle^37^, likely contributes to this NMD blockade. From a viral fitness perspective, our results support the antiviral role of NMD during viral infection, as it promotes the interferon response which limits viral replication (**Figure 4A**, **Figure 6**). This model is in agreement with the widespread observation of NMD inhibition by viruses, likely globally benefiting viral fitness^17–20^. Since NMD is antiviral, tolerating some levels of dsRNA and translation inhibition during infection might even prove beneficial for viruses as a way to inhibit NMD and limit the antiviral cellular responses. However, the discovery of this negative feedback loop involving NMD fails to provide a clear explanation for the proviral role of NMD during some viral infections, such as HCV and HIV^24,25^. One possibility would be that the sensitivity to NMD is influenced by sensitivity to the IFN response; viruses with a low sensitivity to the IFN response would be also lowly sensitive to this negative feedback loop involving NMD and might have evolved further mechanisms to subvert NMD and make it proviral. NMD’s impact likely extend to other cellular processes during viral infection beyond dsRNA sensing, both in host pathways as well as direct effect on the viral replication cycle, and further meticulous studies will be required to decipher each respective contribution to the global output observed. Notably, since the role of NMD during DNA viruses and −ssRNA viruses such as IAV and VSV has been less studied, a better characterization of the role and status of NMD during these infections will be helpful^19^.

**Figure 6.**
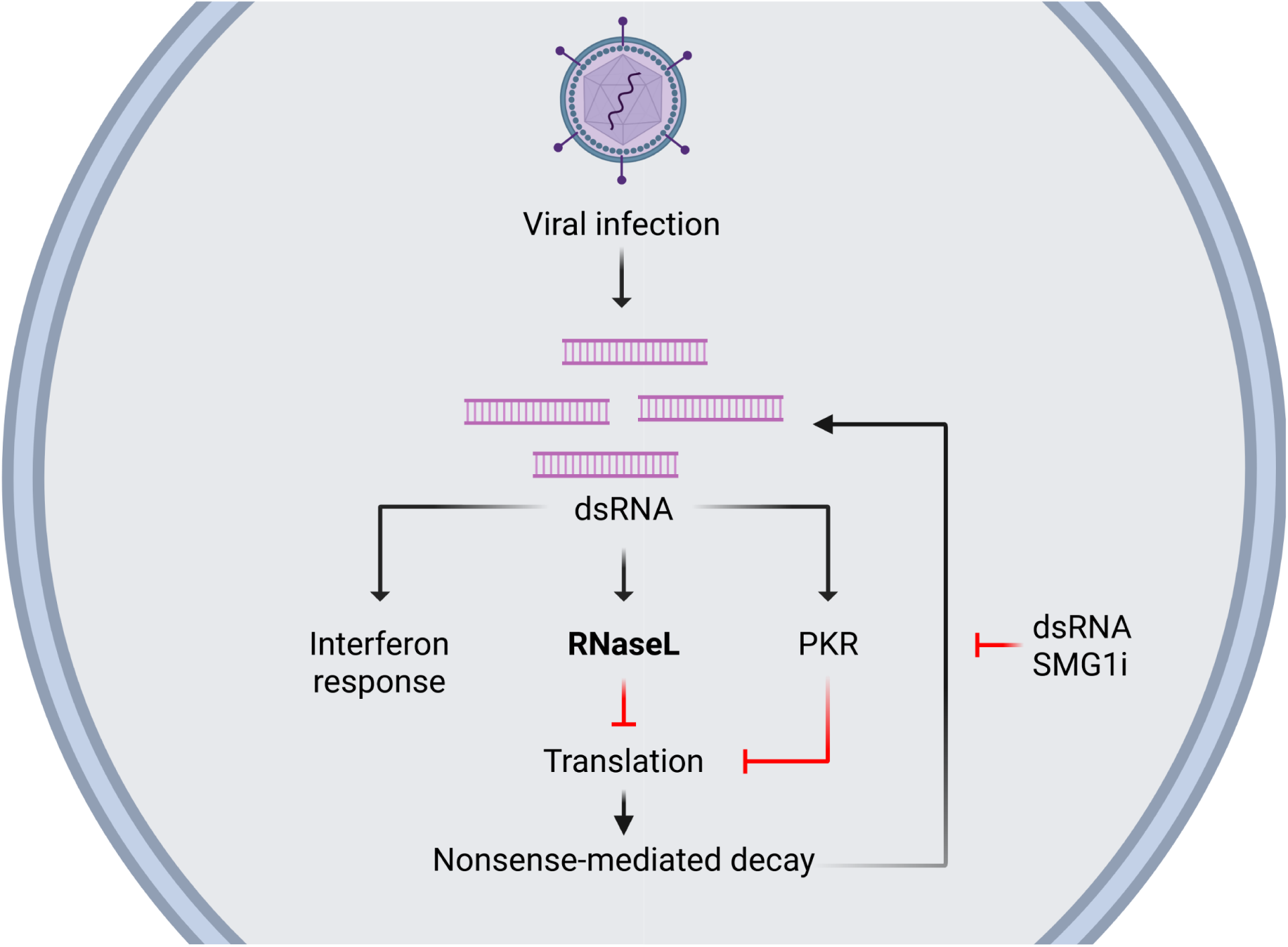
Model of the feedback loop involving NMD during dsRNA sensing. Upon viral infection, dsRNA is produced by most viral infections. This dsRNA, aside from triggering the innate immune sensing pathway and the interferon response, activates RNaseL and PKR, which drastically inhibits translation. In this state of stricking reduction of translation, NMD is also decreased, leading to the accumulation of NMD-decayed isoforms. These isoforms, or the inhibition of NMD itself, feeds back to limit the sensing of dsRNA, in a negative feedback look that will decrease the interferon response as well as PKR and RNaseL activation, and PIC-mediated cell death. Created in BioRender. Lynch, K. (2025) https://BioRender.com/3or17n1.

Interestingly, we don’t observe any effect of SMG1i on IFN-β induction at the highest PIC concentration (**Supplemental Figure S6A**), and RNaseL activation (**Supplemental Figure S10**), contrary to PIC-mediated cell death as well as PKR, for which we do observe an impact of SMG1i at this highest PIC concentration (**Figure 5A, 5B, Supplemental Figure S10**). These results suggest that the former pathways are already being activated by PIC past the linear dose-response range, in opposition to the latter, which are likely less sensitive and require a higher dsRNA load for activation. The need to use systems with no/low NMD inhibition to decipher this feedback loop for some pathways, such as the RNaseL/PKR double-depleted cells and lower levels of PIC, indicate that the inhibitory effect of NMD on these pathways happens at low level of inhibition and quickly reach a maximal effect. Moreover, it also suggests this feedback happens under physiological circumstances to fine-tune and limit the inflammation process, likely contributing with other control mechanisms to the overall prevention of over activation. Limited evidence of the control of inflammation by NMD has been shown in human^38,39^, but some evidence exist in plants^40,41^. Further studies should aim to decipher more broadly the role of NMD in inflammation and immunity.

One remaining question is the exact molecular mechanism explaining this decrease in dsRNA content and sensing upon NMD inhibition. Even though one logical explanation could have involved UPF1, which is an helicase that has been observed to bind viral RNA as well as dsRNA^42–47^, the fact that most of our experiment with SMG1i could be replicated with the knockdown of UPF1 suggest UPF1 is dispensable for this effect. The most likely explanation is that NMD inhibition leads to the activation/increase in expression of a helicase that mediates increased dsRNA unwinding. However, investigations into the most likely culprits, such as SKIV2L^48,49^, DHX9^50^, and EIF4A1/EIF4A2 have proven that these helicases are not involved in mediating this NMD effect on dsRNA sensing (**Supplemental Figure S10**). Further studies should aim the identify the downstream effector of this NMD-mediated dsRNA sensing control.

In conclusion, we have established that NMD is inhibited during dsRNA sensing, and this inhibition limits dsRNA sensing (**Figure 6**). The main functional driver of this dsRNA-induced NMD inhibition is RNaseL, as removal of RNaseL increases the induction of the interferon response without a significant contribution of PKR (**Figure 4A**). This is intriguing as most viruses don’t activate RNaseL^34^. However, it has already been shown that viruses exert direct influence on the NMD machinery^16^. Moreover, we have also shown that in the absence of RNaseL, PKR activation drives at least some inhibition of NMD (**Figure 3A, 3B**). Therefore, we conclude that the precise mechanism by which NMD inhibition is achieved likely varies in a virus-specific manner, and further studies should aim to precisely decipher the status of NMD in viral-infected cells, the respective roles of RNaseL and PKR in these systems, and potential contributions from alternative splicing and other mechanisms to the accumulation of these NMD isoforms. However, our data demonstrates that regardless of the mechanism by which inhibition of NMD is achieved, at least one consequence is the attenuation of further dsRNA sensing. Therefore, we have uncovered a negative feedback loop in innate immune sensing involving NMD that contributes to fine-tuning the cellular response to dsRNA and, likely, viral infection.

## MATERIAL AND METHODS

### Cells, drugs, and viruses

A549 were grown in RPMI medium (Corning, 10-040-CV) supplemented with 10% heat-inactivated fetal bovine serum (FBS) (Gibco, A56697-01), and 2 mM penicillin-streptomycin (Mediatech, MT-30-002-Cl). The generation of OAS1, OAS2, OAS3, RNaseL, and PKR KO A549 cells was described before^34,51^. HEK 293T were grown in DMEM (Corning, 10-013-CV) supplemented with 10% FBS, 2 mM penicillin-streptomycin. L929 were grown in EMEM (ATCC, 30-2023) supplemented with 5% FBS and 2 mM penicillin-streptomycin. All cell lines were maintained at 37°C in a humidified incubator with 5% CO_2_. Cycloheximide (CHX, MilliporeSigma) and homoharringtonine (HHT, Abcam) were diluted in DMSO and used at a final concentration of 100 μg/mL (CHX) or 2 μg/mL (HHT). hSMG-1 inhibitor 11j (SMG1i, MedChemExpress) was diluted in DMSO and used at 3 μM, except where indicated. Okadaic Acid and Calyculin (Millipore Sigma) were diluted in DMSO and used at 500 nM and 50 nM, respectively. Vesicular stomatitis virus harboring eGFP (VSV-GFP) was kindly provided by Dr. John K. Rose and described before^52^. Mayaro virus (MAYV, strain BeH407), Zika virus (ZIKV, strain MR766), and Rift Valley Fever virus (RVFV, strain MP12) were gifts from Dr. Michael Diamond. West Nile virus (WNV, KUNV isolate CH16532) was a generous gift of Dr. Robert Tesh. Chikungunya virus (CHIKV, strain Ross) was provided by Dr. David Weiner (University of Pennsylvania). Influenza A virus (IAV) used in this manuscript is A/WSN/33 (WSN). CHIKV, MAYV, WNV, and ZIKV were all propagated in C6/36 mosquito cells. RVFV and VSV-GFP were propagated in BHK-21 cells. IAV was propagated in MDCK cells. Supernatants were collected, aliquoted and subjected to only a single freeze-thaw. Viral titers were determined by plaque assays on BHK-21, Vero-E6 cells, or MDCK cells. Vaccinia virus (VACV, Copenhagen strain, VC-2) and the ΔE3L mutant^53^ were propagated and titered as previously described^54^. VSV-GFP was UV-inactivated in 1.5 mL tubes, lid opened, 100 μL/tube, under a 254 nm handheld UV lamp (UVP UVG-11, Analytikjena) for 15 min. Controls were infected for longer timepoints alongside the experiment and monitored for GFP expression to validate abrogation of viral replication. Human IFN-β recombinant protein (ThermoFisher, 300-02BC) was used at 1, 10, or 100 units/mL for 5h.

### RNA sequencing and bioinformatic analyses

Public datasets include A549 cells infected with WSN strain of IAV^13^, PIC-stimulated cells^55–58^, and NMD perturbation in HEK 293^59^ and Hela^4^. A549 cells were infected with VSV-GFP at an MOI of 5 or mock infected. RNA was harvested by Trizol at 24 hpi and then analyzed by RNA sequencing using 150 bp paired end reads with the Illumina technology. Raw fastq files from this experiment, as well as raw fastq files downloaded from published datasets, were then trimmed with bbduck 38.79 and aligned to version hg38 of the human genome with STAR 2.7.10a producing BAM files that were indexed with samtools. Next, the bam files were analyzed using MAJIQ 2.4 and batch corrected with moccasin 0.25. Local splicing variations (LSV) were called as significant if the probability of a delta-psi of 10% was greater than 95%. For pie charts, an LSV was considered as NMD-induced if it was significant in at least one NMD dataset. An LSV was considered PIC-induced if it was significant in at least two datasets.

### PAMP, siRNA, and RNA transfection

PAMP were bought from Invivogen: Poly(I:C) HMW (tlrl-pic); Poly(I:C) LMW (tlrl-picw), Poly(A:U) (tlrl-pau), 5’-OH-dsRNA (tlrl-3prnac), 5’-PPP-dsRNA (tlrl-3prna), 5’-PPP-hpRNA (tlrl-hprna) and Poly(dA:dT) (tlrl-patn). 1 μg/mL of Poly(I:C) HMW was used except where indicated; the volume refers to the volume the cells are grown in. For PAMP transfection, 1 μg of the PAMP was transfected using 3 μL of Lipofectamine2000 (ThermoFisher Scientific) in a 12-well plate or scaled accordingly and RNA harvested after 6h. For siRNA transfection, 10 pmoles of siRNA was transfected using 3 μL of RNAiMAX (ThermoFisher Scientific) in 12-well plate or scaled accordingly. For experiment where multiple targets were depleted, 10 pmoles of each siRNA was transfected with 3 μL of RNAiMAX. Cells were incubated for 72 h to allow complete depletion of the target. ON-TARGETplus SMARTpool siRNA were purchased from Horizon Discovery against DHX9 (L-009950-00-0005), PKR (L-003527), RNaseL (L-005032), UPF1 (L-011763), or non-targeting control (D-001810-10). ON-TARGETplus siRNA were purchased from Horizon Discovery against EIF4A1 (J-020178-06), EIF4A2 (J-013758-10), or Non-targeting Control siRNA #2 (D-001810-02). For RNA transfection, 1 μg of trizol-extracted RNA was transfected with 3 μL Lipofectamine MessengerMAX (ThermoFisher Scientific), and RNA harvested after 6h.

### RNA extraction

Total RNA was extracted using TRIzol (Invitrogen) according to the manufacturer’s instructions.

### Radioactive RT-PCR

Low-cycle radioactive RT-PCR were conducted as previously described^60,61^. 250 ng of RNA was reverse transcribed using MMLV reverse transcriptase (ThermoFisher) using the same reverse primer used for the PCR step. Primer sequences are listed in Supplementary Table 1. PCR reactions were carried out for between 20 to 25 cycles depending on the target. Amplicons were resolved on a 5% denaturing polyacrylamide-Urea gel, densitometry measured using a Typhoon PhosphorImager (Amersham Biosciences) and quantitated using Fiji.

### qPCR

The complete list of primers used in this study is available in Supplementary Table 1. Reverse transcription was done using 1 μg of RNA, random hexamers (Invitrogen), and the MMLV reverse transcriptase (Thermo, 28025013). cDNA was diluted to 5 ng/μL following the RT step. Quantitative PCR (qPCR) reactions were prepared in 10 μl in 384-well plates with 5 μL of SYBR qPCR Master Mix (Applied Biosystems, A25742), 2 μL of cDNA (10 ng), 0.4 μL of the primer pair at 10 μM, and 2.6 μL of water. qPCR reactions were carried on a QuantStudio 6 Flex Real-Time PCR System (Applied Biosystems) and data acquired with QuantStudio 12K Flex software version 1.3 (Applied Biosystems). ACTB was used for normalization upon IFN-β treatment; for qPCR on PIC-treated samples, ACTB was unsuitable for normalization as RNaseL activation degrades mRNA^28^. For these samples, we normalized gene expression of the lncRNA MALAT1, which is nuclear and its expression level insensitive to RNaseL activation. For UPF1 depletion, since MALAT1 expression is controlled by UPF1^62^, another non-coding nuclear RNA insensitive to UPF1 depletion and RNaseL activation, 7SK, was used as the housekeeping gene for normalization. For all qPCR experiments, control reactions (no RT) were performed in the absence of cDNA for each primer pair, and confirmed to be negative.

### Protein harvesting and Western-Blot

We first experimentally determined the linearity of all antibodies used in this study for the samples analyzed to allow for an adequate quantification in the linear range. For puromycin-incorporation assay, cells were treated with 10 µg/mL of puromycin for 10 min prior to harvesting, as previously described^63^. Cells were washed with PBS first and then lysed in RIPA buffer (150 mM NaCl, 1% NP-40, 0.5% Sodium Deoxycholate, 0.1% SDS, 50 mM Tris (pH 8.0), cOmplete Mini EDTA-free Protease Inhibitor Cocktail (Millipore Sigma) to 1x), 5 min on ice. For phosphorylated proteins, Halt Protease and Phosphatase Inhibitor Cocktail (ThermoFisher Scientific) was added to the RIPA buffer. DNA was fragmented using sonication at 30% amplitude for 1 min (10 seconds on, 10 seconds off) using a qSonica sonicator. Lysates were then cleaned of debris by centrifugation at 13 000 RPM, 4°C, 10 min. Total protein content was measured using a standard Bradford assay. Samples were diluted in Laemmli buffer, heated at 95°C for 5 min and loaded on 10% or 6% SDS-polyacrylamide gels alongside Precision Plus All Blue prestained protein ladder (Bio-Rad). Gels were transferred onto nitrocellulose membranes, blocked in 5% milk or BSA for 1 h and incubated with primary antibody diluted 1:1000 in blocking buffer overnight at 4°C. The commercial primary antibodies used in this study are the following: Actin (Sigma, A5441), DHX9 (Abcam, ab26271), eIF2α (Cell Signaling Technology, 9722S), p-eIF2α (Cell Signaling Technology, 9721S), EIF4A1 (Cell signaling, 2490), EIF4A2 (Abcam, ab31218), PKR (Cell Signaling Technology, 12297), p-PKR (Abcam, ab32036), puromycin (Millipore Sigma, MABE343), RNaseL (Novus Biologicals, NBP2-80929), SKIV2L (Proteintech, 11462-1-AP), UPF1 (Cell Signaling Technology, 9435), p-UPF1 (Millipore Sigma, 07-1016), Vinculin (Santa Cruz Biotechnology, sc-73614). Blots were washed 2 x 1 min, 2 x 10 min and incubated with Donkey Anti-Mouse or Anti-Rabbit IgG Polyclonal Antibody (IRDye® 800CW, 680RD) diluted 1:10,000 for 1 h at room temperature for fluorescent western-blot, or anti-rabbit HRP-linked secondary antibody (Cell signaling, 7074S, 1:3000) or anti-mouse HRP-linked secondary antibody (Cell signaling, 7076S, 1:5000) for HRP-based imaging. Membranes were washed again 2 x 1 min, 3 x 10 min and and imaged on an Odyssey M Imaging System (LI-COR Biosciences) for fluorescent western-blot. Images were analyzed with Empiria studio version 3.2 (LI-COR). For HRP-based imaging, signal was visualized upon addition of Amersha ECL prime western blotting detection reagent (Cytiva) and subsequent imaging with an X-ray developer.

### Immunofluorescence

A549 cells were seeded in 24-well plates on #1 glass coverslips (ThermoFisher Scientific) and treated as described in other sections. Cells were washed with PBS and fixed for 20 min using 4% paraformaldehyde in PBS (ThermoFisher Scientific). Cells were permeabilized in 0.15% triton X-100 (in PBS) for 5 min and then washed twice with PBS. Blocking was done in 10% goat serum (Gibco, diluted in PBS) for 20 min. Primary antibody against IRF3 (Cell Signaling, 11904, 1:250) and dsRNA (9D5, Absolute Ab, Ab00458-23.0, 1:500) were diluted in blocking buffer and incubated overnight at 4°C. Cells were washed in blocking buffer, three times, for 5 min. Goat anti-rabbit secondary antibody coupled to Alexa Fluor Plus 488 (ThermoFisher Scientific, A32731) was also diluted in blocking buffer (1:1000) and incubated 1 h in the dark. Cells were washed three time in PBS, and nuclei staining was performed using 1 μg/ml Hoechst 33342 in PBS for 15 min at room temperature. Coverslips were mounted on slides with SlowFade Diamond Antifade mounting medium (Invitrogen).

### Microscopy imaging

Immunofluorescence experiments were imaged using a Zeiss LSM 980 confocal unit (50 μm pinhole size) attached to a Zeiss Axio Observer 7 inverted microscope. Images were acquired using a Plan Apochromat 63x/1.40 oil DIC M27 immersion objective, with a final pixel size of 0.071 μm/pixel. Green and Hoechst fluorescence were collected by illuminating the sample with a solid-state visible light laser at 488 nm and 405 nm, respectively. A 32-channel GaAsP-PMT detector was used to detect the emitted signal for Alexa Fluor 488, and a multialkali-PMT detector was used for Hoechst. The images were captured bidirectionally, with a pixel dwell time of 0.55 μs, a 1 AU pinhole and a repeat per line averaging using the mean intensity. ZEN Blue (v. 3.5, Zeiss) was used to acquire the data which was saved in the czi file format.

### Live-cell imaging

Cells were seeded in black glass-bottomed 12-well plates (Cellvis, P12-1.5H-N), transfected with siRNA for 72h, pre-treated with DMSO or SMG1i for 1 h, and transfected with PIC as described before. 30 minutes after PIC transfection, the media was replaced with Sytox Orange containing-media at 100 nM (ThermoFisher Scientific, S34861) and plates were imaged in a Sartorius Incucyte S3 at 30 min intervals. The analysis was done using the Incucyte software module.

### Data analysis and statistical analyses

All statistical analyses were performed using GraphPad Prism (version 10.4.2). Except were stated otherwise, the statistical results presented are ordinary one-way ANOVA using Dunnett’s multiple comparisons test against the control sample; ns, P > 0.05; *P ≤ 0.05; **P ≤ 0.01; ***P ≤ 0.001; ****P ≤ 0.0001. All results presented in this article are mean ± standard deviation.

## ACKNOWLEDGEMENT

We thank Dr. Susan Weiss for providing the A549 KO cell lines. We would like to thank the CDB Microscopy Core (RRID SCR_022373) for providing access to confocal microscopes as well as technical help and training. We also thank Dr. Sara Cherry and Mark Dittmar for providing RNA from infected cells. FUNDING – This work was supported by R35 GM118048 to KWL and a Banting scholarship from the Canadian Institutes of Health Research (CIHR), as well as postdoctoral scholarships from the Fonds de recherche du Québec – Santé (FRQ-S) and from the CIHR to S.B.

## AUTHOR CONTRIBUTIONS

Conceptualization: SB, KWL; Data curation: SB, YR, MBF, JB; Formal analysis: SB, YR, MBF, JB; Funding acquisition: KWL; Investigation: SB, KWL; Methodology: SB, YR, KWL; Project administration: KWL; Resources: BLJ, KWL; Software: MBF; Supervision: SB, BLJ, KWL; Validation: SB, YR; Visualization: SB, MBF; Writing – original draft: SB, KWL; Writing – review & editing: SB, YR, MBF, JB, BLJ, KWL.

## REFERENCES

1. He, F. & Jacobson, A. Nonsense-Mediated mRNA Decay: Degradation of Defective Transcripts Is Only Part of the Story. Annual Review of Genetics 49, 339–366 (2015).

2. Kurosaki, T. et al. A post-translational regulatory switch on UPF1 controls targeted mRNA degradation. Genes Dev. 28, 1900–1916 (2014).

3. Yamashita, A., Ohnishi, T., Kashima, I., Taya, Y. & Ohno, S. Human SMG-1, a novel phosphatidylinositol 3-kinase-related protein kinase, associates with components of the mRNA surveillance complex and is involved in the regulation of nonsense-mediated mRNA decay. Genes Dev. 15, 2215–2228 (2001).

4. Colombo, M., Karousis, E. D., Bourquin, J., Bruggmann, R. & Mühlemann, O. Transcriptome-wide identification of NMD-targeted human mRNAs reveals extensive redundancy between SMG6- and SMG7-mediated degradation pathways. RNA 23, 189–201 (2017).

5. Muñoz, O., Lore, M. & Jagannathan, S. The long and short of EJC-independent nonsense-mediated RNA decay. Biochemical Society Transactions BST20221131 (2023) doi:10.1042/BST20221131.

6. Kurosaki, T., Popp, M. W. & Maquat, L. E. Quality and quantity control of gene expression by nonsense-mediated mRNA decay. Nat Rev Mol Cell Biol 20, 406–420 (2019).

7. Saltzman, A. L. et al. Regulation of Multiple Core Spliceosomal Proteins by Alternative Splicing-Coupled Nonsense-Mediated mRNA Decay. Molecular and Cellular Biology 28, 4320–4330 (2008).

8. Frankiw, L., Mann, M., Li, G., Joglekar, A. & Baltimore, D. Alternative splicing coupled with transcript degradation modulates OAS1g antiviral activity. RNA 26, 126–136 (2020).

9. Ni, J. Z. et al. Ultraconserved elements are associated with homeostatic control of splicing regulators by alternative splicing and nonsense-mediated decay. Genes Dev. 21, 708–718 (2007).

10. Boudreault, S. et al. Global Profiling of the Cellular Alternative RNA Splicing Landscape during Virus-Host Interactions. PLoS One 11, e0161914 (2016).

11. Boudreault, S. et al. Reovirus μ2 protein modulates host cell alternative splicing by reducing protein levels of U5 snRNP core components. Nucleic Acids Res. 50, 5263–5281 (2022).

12. Boudreault, S., Roy, P., Lemay, G. & Bisaillon, M. Viral modulation of cellular RNA alternative splicing: A new key player in virus–host interactions? Wiley Interdiscip. Rev. RNA 10, e1543 (2019).

13. Thompson, M. G. et al. Viral-induced alternative splicing of host genes promotes influenza replication. eLife 9, e55500 (2020).

14. Mann, J. T., Riley, B. A. & Baker, S. F. All differential on the splicing front: Host alternative splicing alters the landscape of virus-host conflict. Seminars in Cell & Developmental Biology 146, 40–56 (2023).

15. Balistreri, G. et al. The Host Nonsense-Mediated mRNA Decay Pathway Restricts Mammalian RNA Virus Replication. Cell Host & Microbe 16, 403– 411 (2014).

16. Li, M. et al. Identification of antiviral roles for the exon–junction complex and nonsense-mediated decay in flaviviral infection. Nat Microbiol 4, 985–995 (2019).

17. Ahmed, M. R. & Du, Z. Molecular Interaction of Nonsense-Mediated mRNA Decay with Viruses. Viruses 15, 816 (2023).

18. Popp, M. W.-L., Cho, H. & Maquat, L. E. Viral subversion of nonsense-mediated mRNA decay. RNA 26, 1509–1518 (2020).

19. van der Klugt, T. & van Gent, M. The dynamic interactions between virus infections and nonsense-mediated decay. Human Molecular Genetics ddae151 (2025) doi:10.1093/hmg/ddae151.

20. Leon, K. & Ott, M. An ‘Arms Race’ between the Nonsense-mediated mRNA Decay Pathway and Viral Infections. Seminars in Cell & Developmental Biology 111, 101–107 (2021).

21. Contu, L., Balistreri, G., Domanski, M., Uldry, A.-C. & Mühlemann, O. Characterisation of the Semliki Forest Virus-host cell interactome reveals the viral capsid protein as an inhibitor of nonsense-mediated mRNA decay. PLOS Pathogens 17, e1009603 (2021).

22. Mallick, M. et al. Modulation of UPF1 catalytic activity upon interaction of SARS-CoV-2 Nucleocapsid protein with factors involved in nonsense mediated-mRNA decay. Nucleic Acids Research 52, 13325–13339 (2024).

23. Nuccetelli, V. et al. The SARS-CoV-2 nucleocapsid protein interferes with the full enzymatic activation of UPF1 and its interaction with UPF2. Nucleic Acids Research 53, gkaf010 (2025).

24. Ramage, H. R. et al. A Combined Proteomics/Genomics Approach Links Hepatitis C Virus Infection with Nonsense-Mediated mRNA Decay. Molecular Cell 57, 329–340 (2015).

25. Serquiña, A. K. P. et al. UPF1 Is Crucial for the Infectivity of Human Immunodeficiency Virus Type 1 Progeny Virions. Journal of Virology 87, 8853–8861 (2013).

26. Tran, G. V. Q. et al. Nonsense-mediated mRNA decay does not restrict influenza A virus propagation. Cellular Microbiology 23, e13323 (2021).

27. Fensterl, V., Chattopadhyay, S. & Sen, G. C. No Love Lost Between Viruses and Interferons. Annu. Rev. Virol. 2, 549–572 (2015).

28. Karasik, A. & Guydosh, N. R. The Unusual Role of Ribonuclease L in Innate Immunity. WIREs RNA 15, e1878 (2024).

29. Gal-Ben-Ari, S., Barrera, I., Ehrlich, M. & Rosenblum, K. PKR: A Kinase to Remember. Front. Mol. Neurosci. 11, (2019).

30. Bresson, S., Sani, E., Armatowska, A., Dixon, C. & Tollervey, D. The transcriptional and translational landscape of HCoV-OC43 infection. PLOS Pathogens 21, e1012831 (2025).

31. Gopalsamy, A. et al. Identification of pyrimidine derivatives as hSMG-1 inhibitors. Bioorganic & Medicinal Chemistry Letters 22, 6636–6641 (2012).

32. Jensen, S. & Thomsen, A. R. Sensing of RNA Viruses: a Review of Innate Immune Receptors Involved in Recognizing RNA Virus Invasion. J Virol 86, 2900–2910 (2012).

33. Ishihara, H. et al. Calyculin A and okadaic acid: Inhibitors of protein phosphatase activity. Biochemical and Biophysical Research Communications 159, 871–877 (1989).

34. Li, Y. et al. Activation of RNase L is dependent on OAS3 expression during infection with diverse human viruses. Proceedings of the National Academy of Sciences 113, 2241–2246 (2016).

35. Weber, F., Wagner, V., Rasmussen, S. B., Hartmann, R. & Paludan, S. R. Double-Stranded RNA Is Produced by Positive-Strand RNA Viruses and DNA Viruses but Not in Detectable Amounts by Negative-Strand RNA Viruses. Journal of Virology 80, 5059–5064 (2006).

36. Son, K.-N., Liang, Z. & Lipton, H. L. Double-Stranded RNA Is Detected by Immunofluorescence Analysis in RNA and DNA Virus Infections, Including Those by Negative-Stranded RNA Viruses. Journal of Virology 89, 9383– 9392 (2015).

37. Rozman, B., Fisher, T. & Stern-Ginossar, N. Translation—A tug of war during viral infection. Molecular Cell 83, 481–495 (2023).

38. Oliveira, V. et al. A Protective Role for the Human SMG-1 Kinase against Tumor Necrosis Factor-α-induced Apoptosis *. Journal of Biological Chemistry 283, 13174–13184 (2008).

39. Johnson, J. L. et al. Inhibition of Upf2-Dependent Nonsense-Mediated Decay Leads to Behavioral and Neurophysiological Abnormalities by Activating the Immune Response. Neuron 104, 665–679.e8 (2019).

40. Gloggnitzer, J. et al. Nonsense-Mediated mRNA Decay Modulates Immune Receptor Levels to Regulate Plant Antibacterial Defense. Cell Host & Microbe 16, 376–390 (2014).

41. Nasim, Z., Karim, N., Blilou, I. & Ahn, J. H. NMD-mediated posttranscriptional regulation fine-tunes the NLR-WRKY regulatory module to modulate bacterial defense response. Plant Sci 356, 112528 (2025).

42. Lee, S. et al. The SARS-CoV-2 RNA interactome. Molecular Cell 81, 2838–2850.e6 (2021).

43. Schmidt, N. et al. The SARS-CoV-2 RNA–protein interactome in infected human cells. Nat Microbiol 6, 339–353 (2021).

44. Lim, J. et al. Cellular dsRNA interactome captured by K1 antibody reveals the regulatory map of exogenous RNA sensing. Commun Biol 8, 1–12 (2025).

45. Pennemann, F. L. et al. Cross-species analysis of viral nucleic acid interacting proteins identifies TAOKs as innate immune regulators. Nat Commun 12, 7009 (2021).

46. Fischer, J. W., Busa, V. F., Shao, Y. & Leung, A. K. L. Structure-Mediated RNA Decay by UPF1 and G3BP1. Molecular Cell 78, 70–84.e6 (2020).

47. Girardi, E. et al. Proteomics-based determination of double-stranded RNA interactome reveals known and new factors involved in Sindbis virus infection. RNA 29, 361–375 (2023).

48. Eckard, S. C. et al. The SKIV2L RNA exosome limits activation of the RIG-I-like receptors. Nat Immunol 15, 839–845 (2014).

49. Yang, K., Dong, B., Asthana, A., Silverman, R. H. & Yan, N. RNA helicase SKIV2L limits antiviral defense and autoinflammation elicited by the OAS-RNase L pathway. The EMBO Journal 43, 3876–3894 (2024).

50. Zhang, Z., Yuan, B., Lu, N., Facchinetti, V. & Liu, Y.-J. DHX9 Pairs with IPS-1 To Sense Double-Stranded RNA in Myeloid Dendritic Cells. The Journal of Immunology 187, 4501–4508 (2011).

51. Li, Y. et al. Ribonuclease L mediates the cell-lethal phenotype of double-stranded RNA editing enzyme ADAR1 deficiency in a human cell line. eLife 6, e25687 (2017).

52. Ramsburg, E. et al. A Vesicular Stomatitis Virus Recombinant Expressing Granulocyte-Macrophage Colony-Stimulating Factor Induces Enhanced T-Cell Responses and Is Highly Attenuated for Replication in Animals. Journal of Virology 79, 15043–15053 (2005).

53. Beattie, E. et al. Reversal of the interferon-sensitive phenotype of a vaccinia virus lacking E3L by expression of the reovirus S4 gene. Journal of Virology 69, 499–505 (1995).

54. Tartaglia, J. et al. NYVAC: A highly attenuated strain of vaccinia virus. Virology 188, 217–232 (1992).

55. Burke, J. M. et al. RNase L activation in the cytoplasm induces aberrant processing of mRNAs in the nucleus. PLOS Pathogens 18, e1010930 (2022).

56. Karasik, A., Lorenzi, H. A., DePass, A. V. & Guydosh, N. R. Endonucleolytic RNA cleavage drives changes in gene expression during the innate immune response. Cell Reports 43, 114287 (2024).

57. Petit, M. J. et al. Nuclear dengue virus NS5 antagonizes expression of PAF1-dependent immune response genes. PLOS Pathogens 17, e1010100 (2021).

58. Kidwell, A. et al. Translation Rescue by Targeting Ppp1r15a through Its Upstream Open Reading Frame in Sepsis-Induced Acute Kidney Injury in a Murine Model. J Am Soc Nephrol 34, 220–240 (2023).

59. Britto-Borges, T., Gehring, N. H., Boehm, V. & Dieterich, C. NMDtxDB: data-driven identification and annotation of human NMD target transcripts. RNA 30, 1277–1291 (2024).

60. Ip, J. Y. et al. Global analysis of alternative splicing during T-cell activation. RNA 13, 563–572 (2007).

61. Rothrock, C., Cannon, B., Hahm, B. & Lynch, K. W. A Conserved Signal-Responsive Sequence Mediates Activation-Induced Alternative Splicing of CD45. Molecular Cell 12, 1317–1324 (2003).

62. Li, L. et al. The Human RNA Surveillance Factor UPF1 Modulates Gastric Cancer Progression by Targeting Long Non-Coding RNA MALAT1. Cellular Physiology and Biochemistry 42, 2194–2206 (2017).

63. Schmidt, E. K., Clavarino, G., Ceppi, M. & Pierre, P. SUnSET, a nonradioactive method to monitor protein synthesis. Nat Methods 6, 275–277 (2009).

