## Supplementary Tables and Figures for "Nonsense-mediated decay controls a negative feedback loop in innate immune sensing"

Table S1. Primers used in this study.

| Target | Assay | Forward primer | Reverse primer |
|---|---|---|---|
| EIF4A2 | RT-PCR | GGGATTGATGTGCAACAAGTG | CCACACCTTTCCTCCCAAATCG |
| CDKN2AIP | RT-PCR | CAAAGTGACAGATGCTCCAACC | TGGCAGAGGTTTTTGCGTGATC |
| IFN-β | qPCR | GCTTCTCCACTACAGCTCTTTC | CAGTATTCAAGCCTCCCATTCA |
| IFIT1 | qPCR | GCCTTGCTGAAGTGTGGAGGAA | ATCCAGGCGATAGGCAGAGATC |
| IFIT2 | qPCR | CACTGCAACCATGAGTGAGAA | AGTAGGTTGCACATTGTGGCT |
| Actin | qPCR (housekeeping) | CCCTGGCATTGCCGACAGG | GCCGATCCACACGGAGTACT |
| Malat1 | qPCR (housekeeping) | GGTGATGCGAGTTGTTCTCCGT | CGGTTTCCTCAAGCTCCGCC |
| 7SK | qPCR (housekeeping) | GAGGGCGATCTGGCTGCGACAT | ACATGGAGCGGTGAGGGAGGAA |

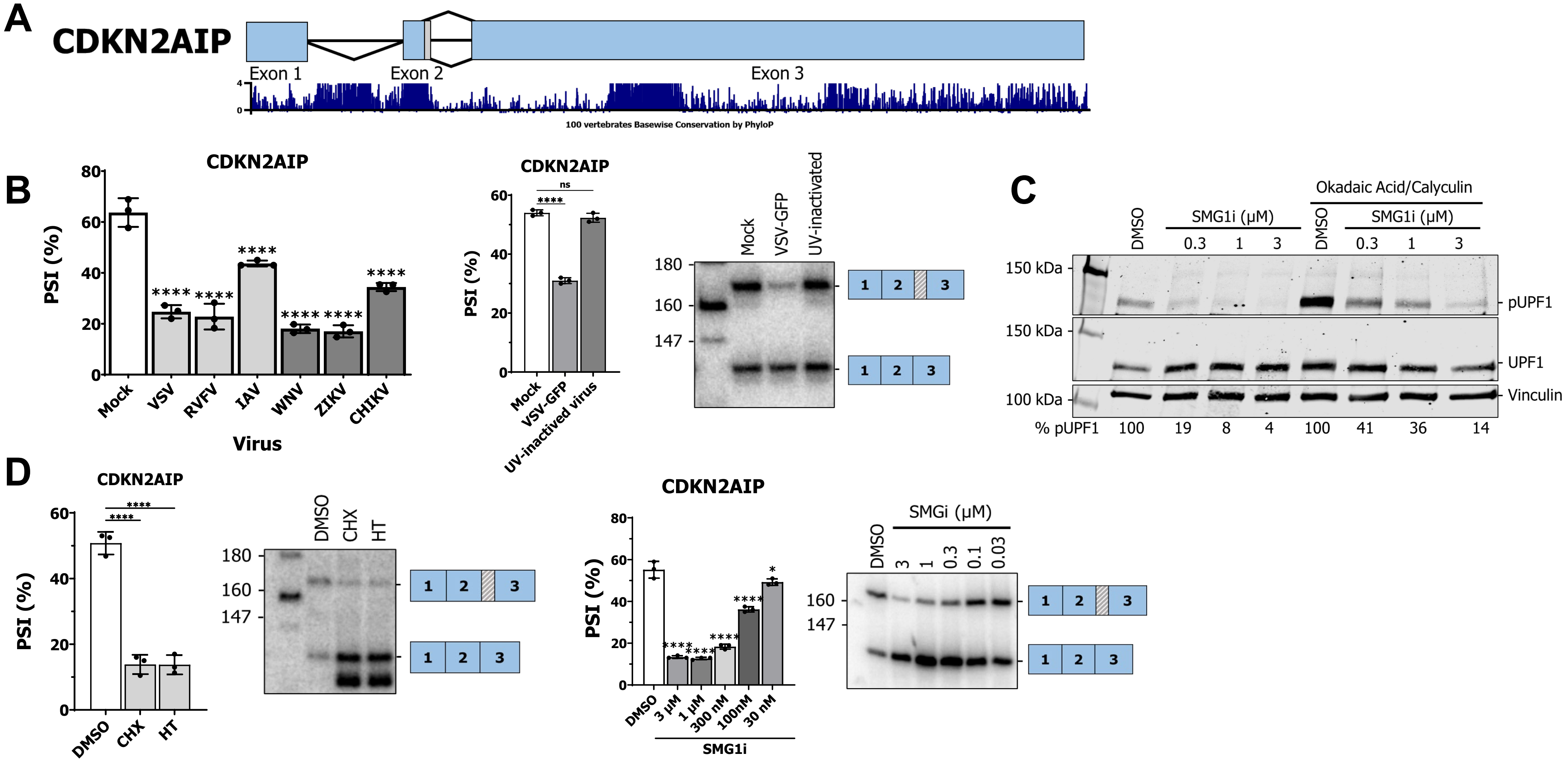

**Figure S1: CDKN2AIP alternative 5′-SS is regulated by viral infection and triggers NMD.** (A) Schematic of the alternative 5′-SS in CDKN2AIP. Conservation values across 100 vertebrates is shown underneath. (B) Left, impact of viral infection with prototypical -ssRNA and +ssRNA viruses on CDKN2AIP 5′-SS usage. Right, impact of VSV and UV-inactivated VSV on CDKN2AIP 5′-SS usage. (C) Western blot of pUPF1, UPF1, and vinculin (loading control) in A549 cells following 7h treatment with SMG1i at 0.3, 1, and 3 μM. 50 nM calyculin and 500 nM okadaic acid were added 5 minutes prior to harvesting to prevent dephospholyration of UPF1 and increase the signal, where indicated. (D) Left, CDKN2AIP 5′-SS usage is regulated by translation inhibition; right, CDKN2AIP 5′-SS usage is regulated by NMD.

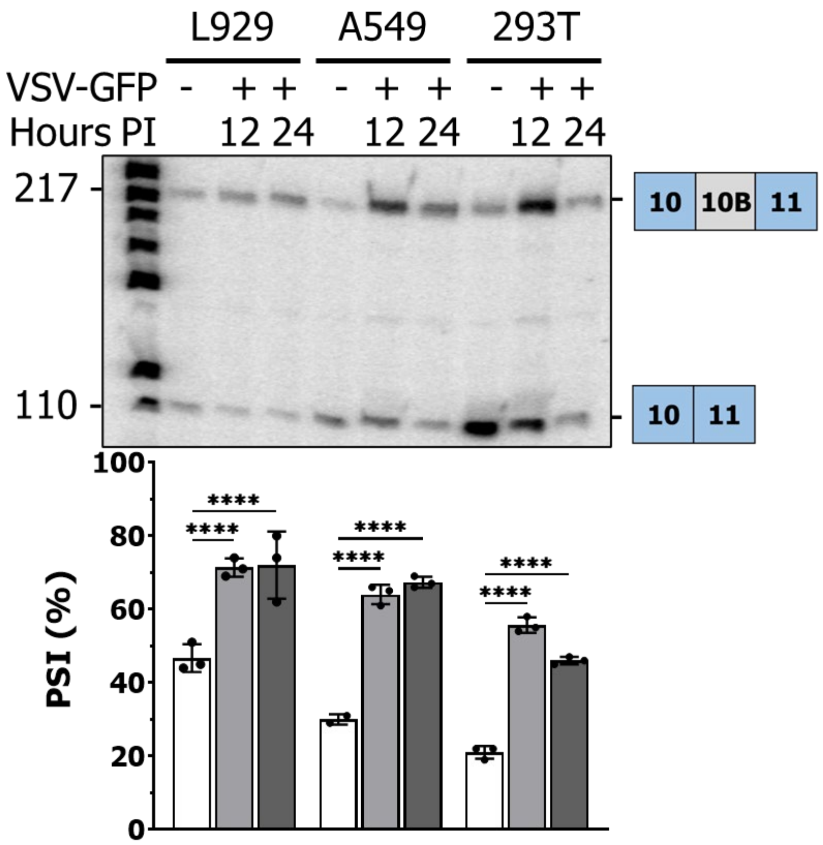


**Figure S2: Impact of VSV-GFP infection on EIF4A2 exon 10B splicing in L929, A549, and HEK 293T**. Cells were infected at an MOI of 5 and harvested at 12 h or 24 h post-infection. Mock-infected controls were harvested at 12h.

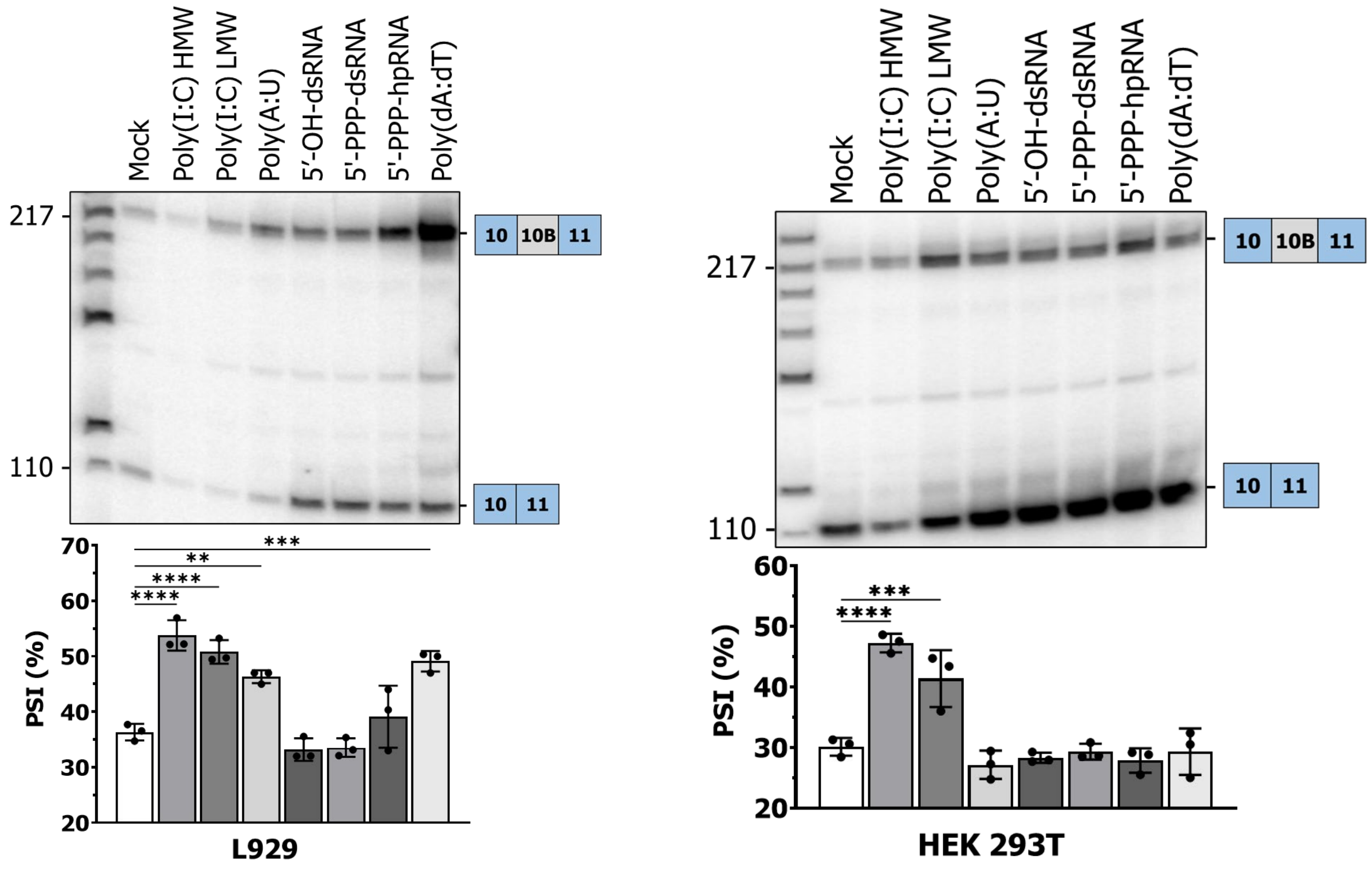


**Figure S3: Monitoring of EIF4A2 exon 10B inclusion following transfection of different PAMP.** Seven different PAMP (1 µg/mL of each) were transfected for 6h. Left, L929 mouse fibroblasts; right, HEK 293T.

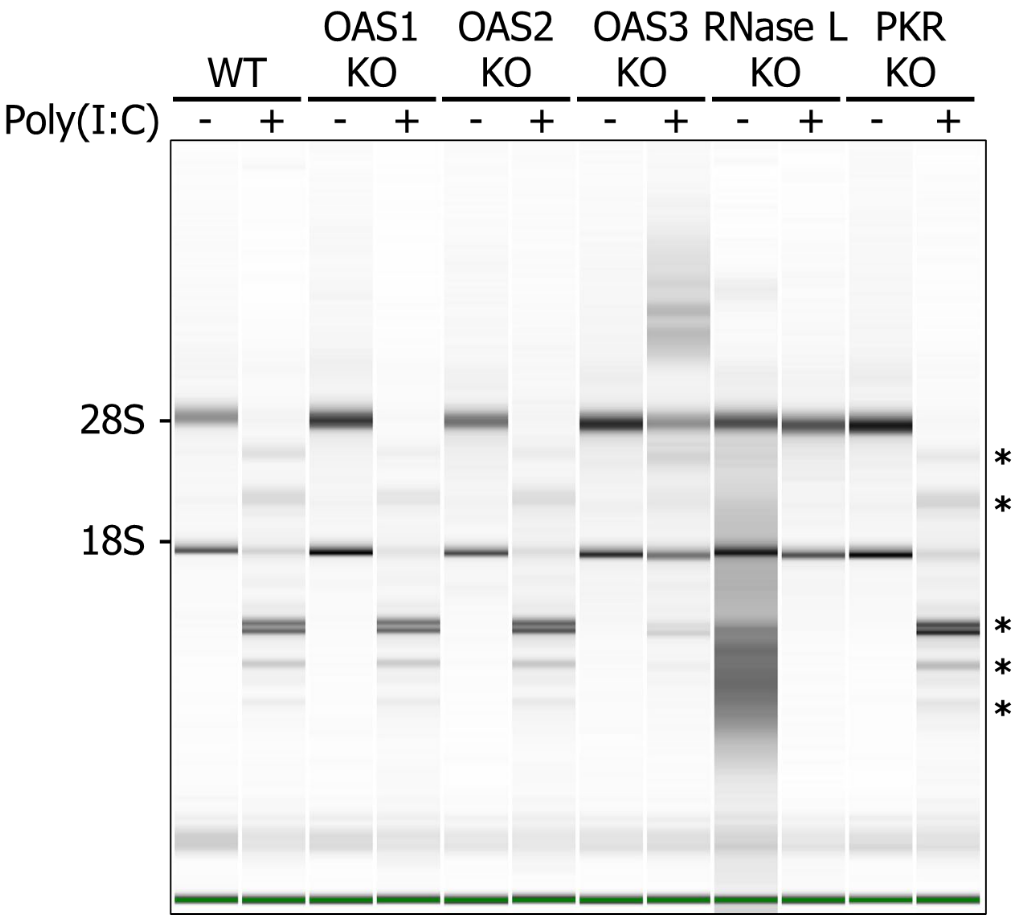


**Figure S4: Validation of RNaseL activation in the different KO cells.** Cells were transfected with 1 µg/mL of PIC for 6 h prior to RNA harvesting and analysis on a Bioanalyzer. Asterisks denote the cleavage products of ribosomal RNA typical of RNaseL activation.

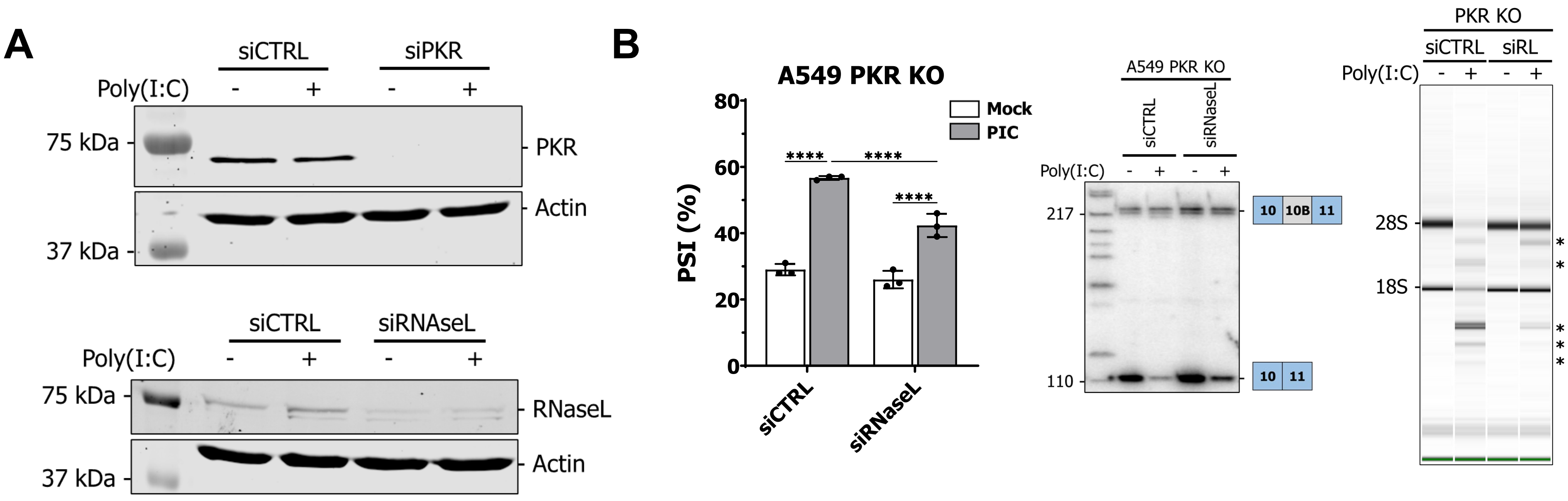


**Figure S5: Impact of RNaseL depletion on EIF4A2 exon 10B inclusion in PKR KO cells.** (A) Top, validation of PKR depletion using siRNA. Bottom, validation of RNaseL depletion using siRNA. (B) Left, impact of RNaseL depletion on EIF4A2 exon 10B inclusion in PKR KO cells. A549 PKR KO cells were transfected with siRNA for 72h, mock transfected or transfected with 1 µg/mL of PIC, RNA harvested after 6 h and subjected to RT-PCR for EIF4A2 exon 10B inclusion. Right, RNA was run on a Bioanalyzer to monitor RNaseL activation. An incomplete reduction of RNaseL activation can be observed in PKR KO treated with siRNase L (siRL). Two-way ANOVA with uncorrected Fisher's LSD multiple comparisons test.

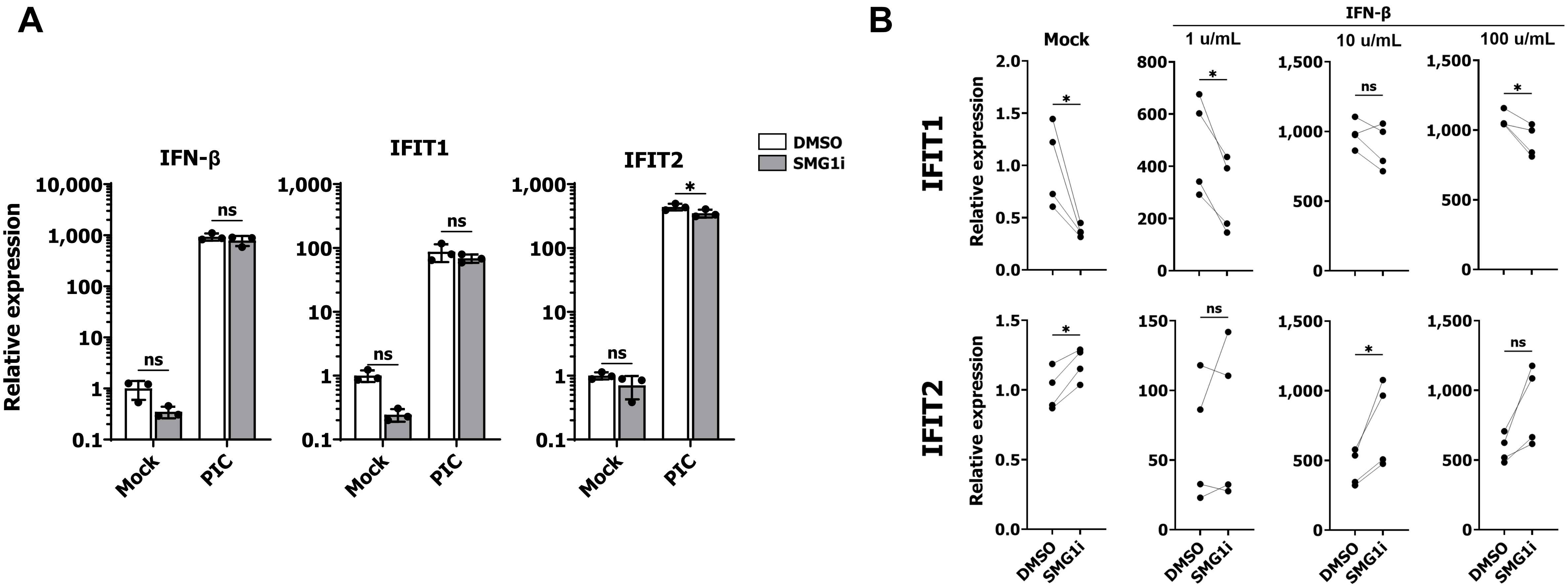


**Figure S6: Impact of NMD inhibition using SMG1i on IFN-β and ISG induction.** (A) A549 cells were pre-treated with SMG1i (3 μM) or DMSO for 1h, transfected with 1 μg/mL of PIC for 6h prior to RNA harvesting and qPCR for IFN-β and IFIT1/2. Two-way ANOVA with uncorrected Fisher's LSD multiple comparisons test. (B) A549 cells were pre-treated with SMG1i (3 μM) or DMSO for 1 h, mock-treated or treated with 1, 10, or 100 u/mL of IFN-β for 5h prior to RNA harvesting and qPCR for IFIT1/2. Paired two-tailed Student's t-test; samples from the same biological replicate were paired together.

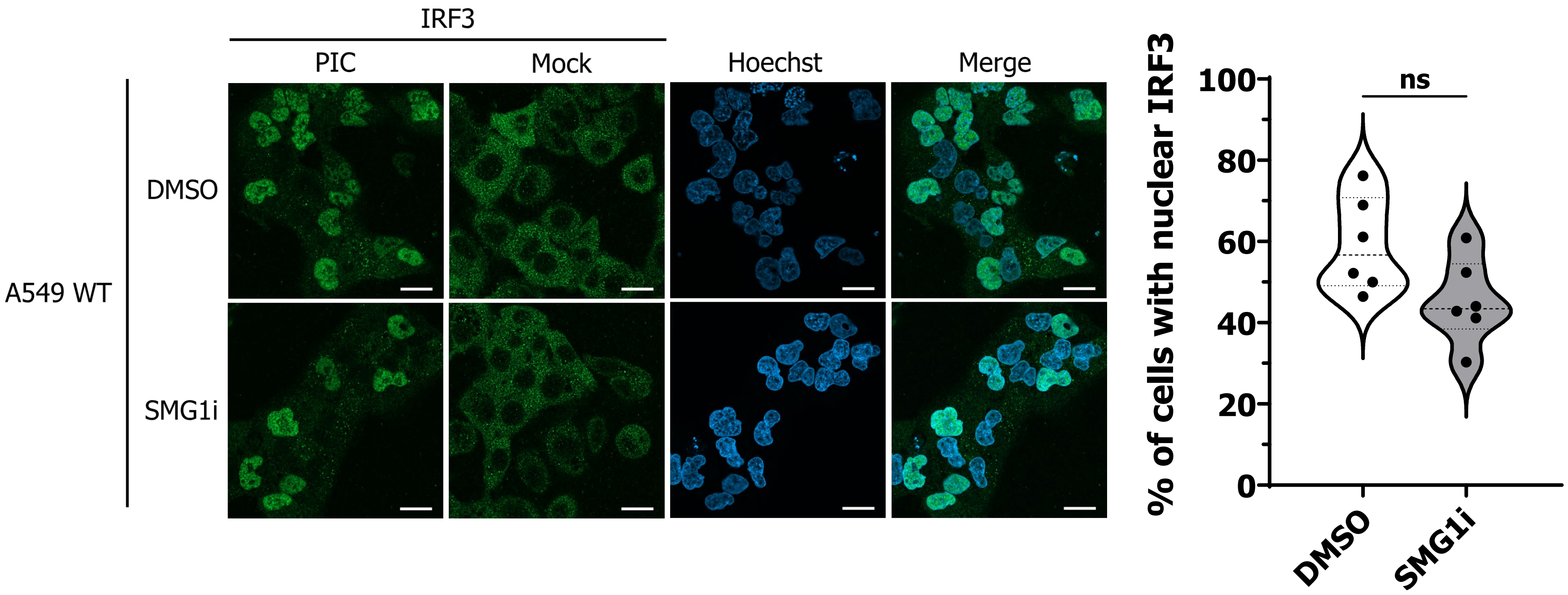


**Figure S7: Impact of SMG1i on IRF3 translocation upon PIC transection.** Left, IRF3 immunofluorescence of DMSO or SMG1i-treated cells transfected with PIC or mock-transfected. WT A549 were transfected with siCTRL for 72h, pre-treated with DMSO or SMG1i (3 µM) for 1 h and transfected with 1 µg/mL of PIC or mock-transfected prior to IF. Scale bar, 20 µm. Right, quantification of the number of cells with IRF3 nuclear localization upon PIC transfection. 6 fields were counted for each condition each containing between 14 and 36 cells. Unpaired two-tailed Student's t-test.

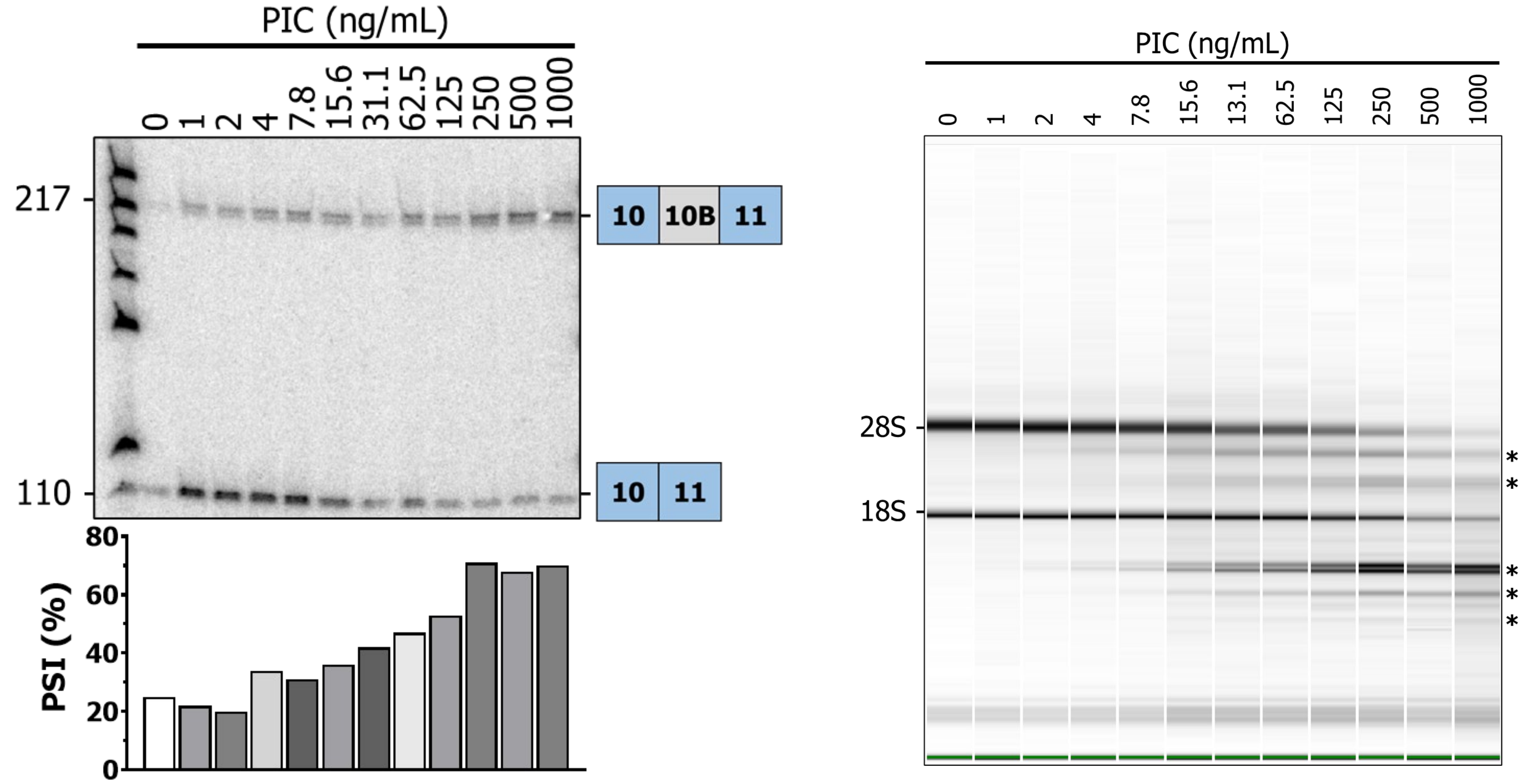


**Figure S8: Impact of different PIC concentrations on NMD inhibition and RNaseL activation.** Left, monitoring of EIF4A2 exon 10B inclusion following transfection of increasing quantity of PIC. Between 0 ng/mL and 1000 ng/mL of PIC was transfected in A549 for 6h prior to RNA harvesting and RT-PCR for EIF4A2 exon 10B. Right, the same RNA were run on a Bioanalyzer to monitor RNaseL activation. Asterisks denote the cleavage products of ribosomal RNA typical of RNaseL activation.

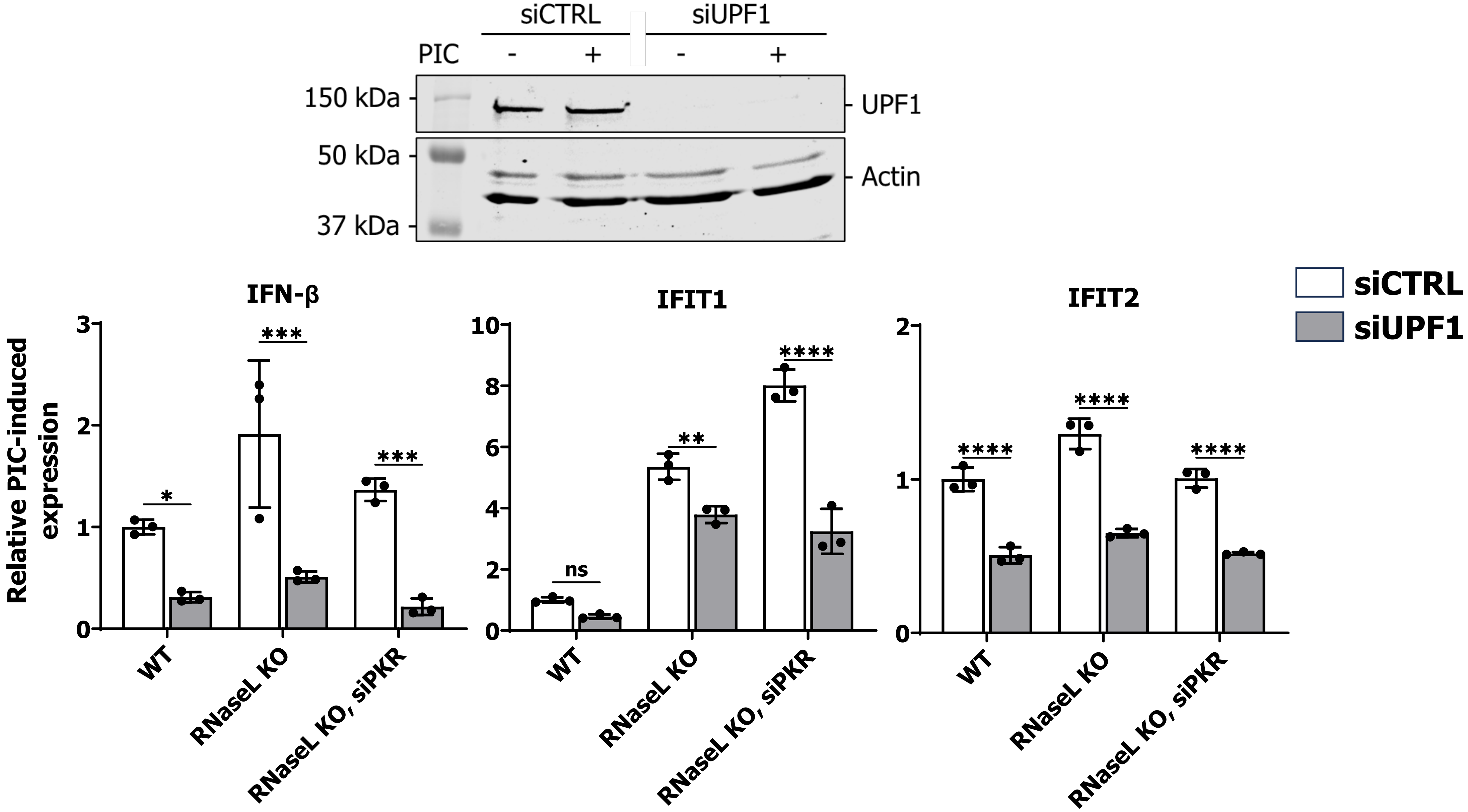


**Figure S9: Impact of siUPF1 on the IFN response.** Top, validation of UPF1 depletion using siRNA. Bottom, A549 WT, RNaseL KO as well as RNaseL KO transfected with siRNA against PKR were transfected with siCTRL or siUPF1 for 72h, transfected with 1 μg/mL of PIC for 6h, and RNA harvested. Two-way ANOVA with uncorrected Fisher's LSD multiple comparisons test.

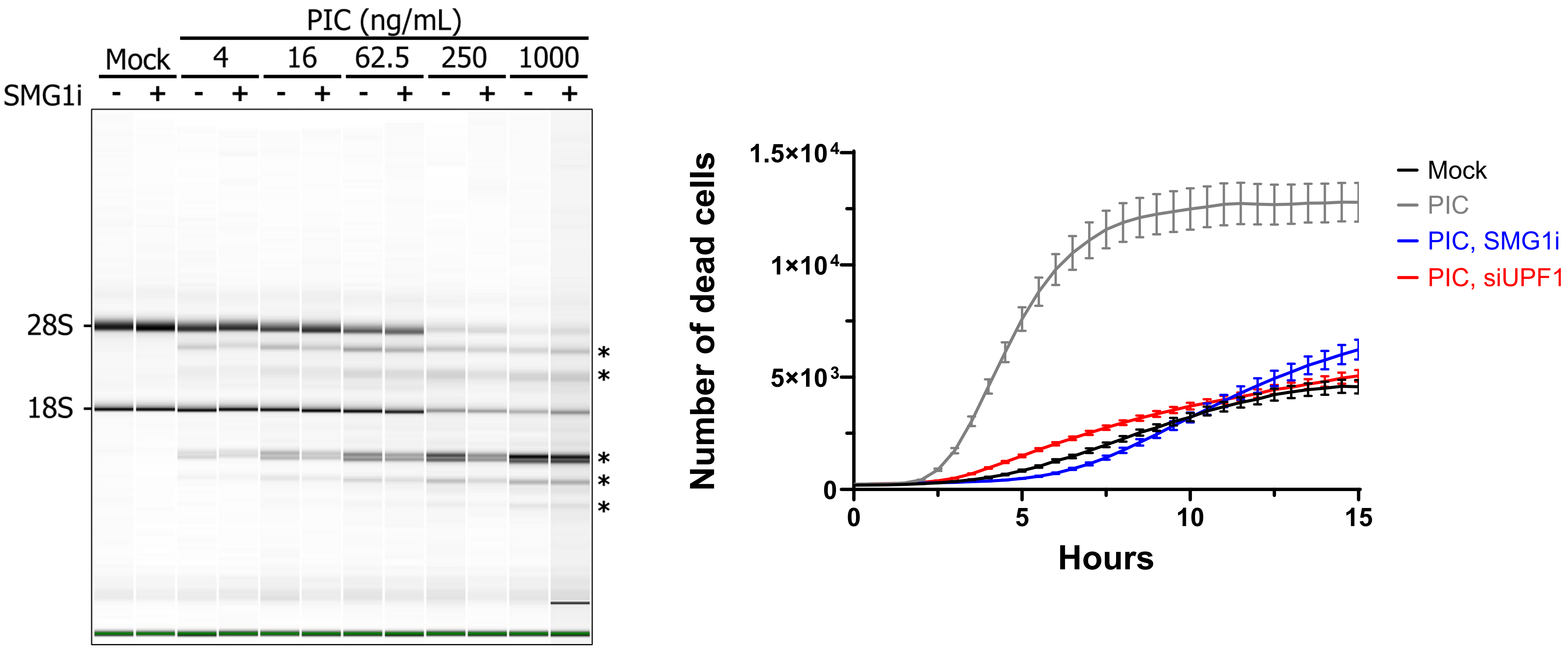


**Figure S10: Impact of NMD inhibition on RNaseL activation and PIC-mediated cell death.** Left, A549 cells were pre-treated for 1 h with DMSO or SMG1i (3 µM) prior to transfection of different concentration of PIC for 6 h; RNA was harvested and run on a Bioanalyzer. Asterisks denote the cleavage products of ribosomal RNA typical of RNaseL activation. Right, live-cell imaging of A549 cells upon transfection of 1 µg/mL of PIC in the presence or absence of 3 µM of SMG1i. Cells were transected with siCTRL or siUPF1 for 72h, transfected with 1 µg/mL of PIC or mock transfected, and monitored with an Incucyte. The SMG1i conditions was pre-treated for 1h before PIC transfection. 30 minutes after PIC transfected, sytox-orange containing medium was added; the number of dead cells correspond to Sytox-Orange positive cells.

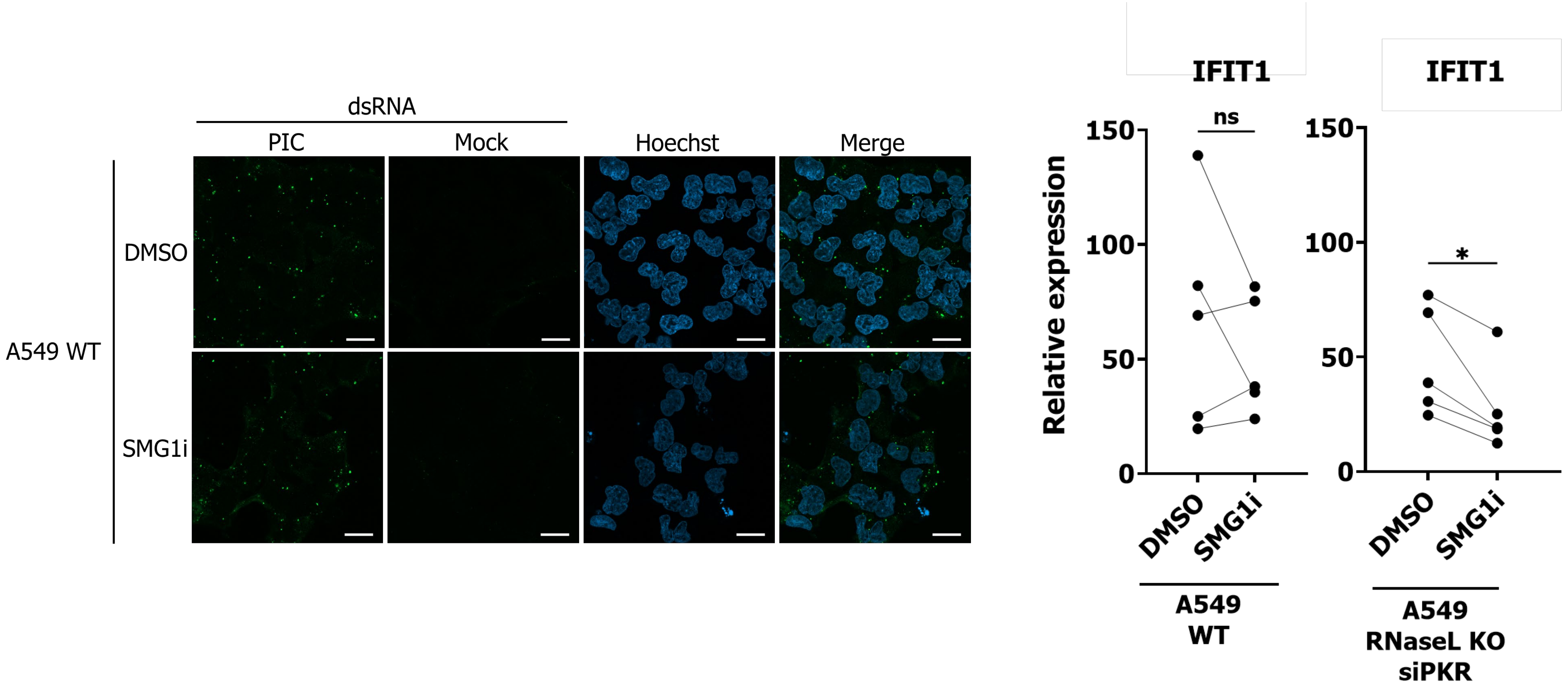


**Figure S11: NMD controls dsRNA load.** Left, immunofluorescence for dsRNA using 9D5 of DMSO or SMG1i-treated cells transfected with PIC in WT A549. Cells were transfected with siCTRL for 72h, pre-treated with DMSO or SMG1i (3 µM) for 1h and transfected with 1 µg/mL of PIC prior to IF using 9D5. Scale bar, 20 µm. Right, RNA harvested from PIC-transfected A549 (treated or not with SMG1i) were transfected in A549 and induction of IFIT1 was measured using qPCR. Expression was normalized against a mock-transfected sample; paired two-tailed Student's t-test (RNA with or without SMG1i from the same experiment).

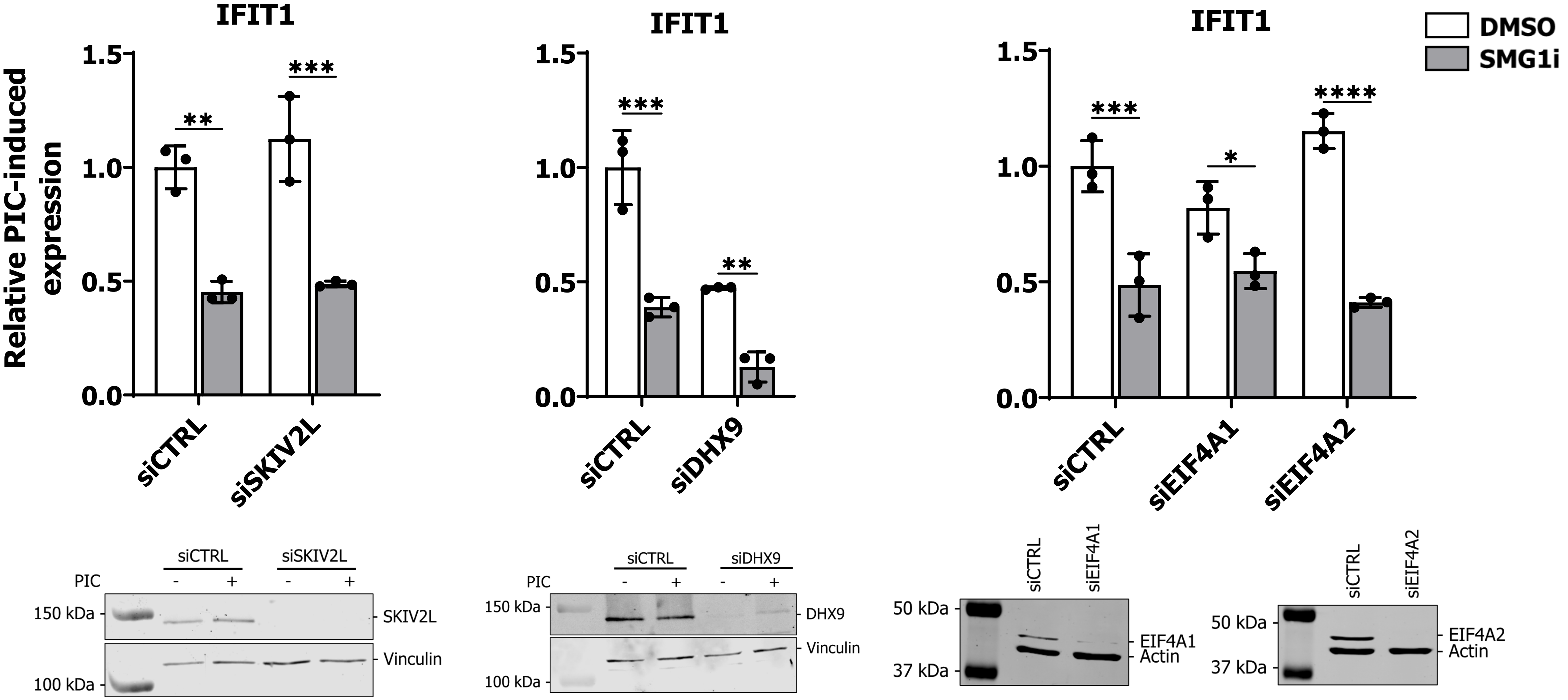


**Figure S12: Impact of helicase depletion on NMD-mediated decrease in the IFN response.** Top, A549 RNaseL KO were transfected with siRNA against PKR, as well as siRNA CTRL or against the different helicases, for 72h. Cells were then pre-treated with DMSO or SMG1i (3 µM) for 1h and transfected with 1 µg/mL of PIC and RNA harvested at 6h for qPCR against IFIT1. Ordinary two-way ANOVA using Tukey's multiple comparisons test. Bottom, validation of SKIV2L, DHX9, EIF4A1, and EIF4A2 depletion using siRNA.
